# Three Reagents for in-Solution Enrichment of Ancient Human DNA at More than a Million SNPs

**DOI:** 10.1101/2022.01.13.476259

**Authors:** Nadin Rohland, Swapan Mallick, Matthew Mah, Robert Maier, Nick Patterson, David Reich

**Author notes:** To whom correspondence should be addressed: N.R., S.M., D.R. Contributed equally: N.R. and S.M.

## Abstract

In-solution enrichment for hundreds of thousands of single nucleotide polymorphisms (SNPs) has been the source of >70% of all genome-scale ancient human DNA data published to date. This approach has made it possible to generate data for one to two orders of magnitude lower cost than random shotgun sequencing, making it economical to study ancient samples with low proportions of human DNA, and increasing the rate of conversion of sampled remains into working data thereby facilitating ethical stewardship of human remains. So far, nearly all ancient DNA data obtained using in-solution enrichment has been generated using a set of bait sequences targeting about 1.24 million SNPs (the ‘1240k reagent’). These sequences were published in 2015, but synthesis of the reagent has been cost-effective for only a few laboratories. In 2021, two companies made available reagents that target the same core set of SNPs along with supplementary content. Here, we test the properties of the three reagents on a common set of 27 ancient DNA libraries across a range of richness of DNA content and percentages of human molecules. All three reagents are highly effective at enriching many hundreds of thousands of SNPs. For all three reagents and a wide range of conditions, one round of enrichment produces data that is as useful as two rounds when tens of millions of sequences are read out as is typical for such experiments. In our testing, the “Twist Ancient DNA” reagent produces the highest coverages, greatest uniformity on targeted positions, and almost no bias toward enriching one allele more than another relative to shotgun sequencing. Allelic bias in 1240k enrichment has made it challenging to carry out joint analysis of these data with shotgun data, creating a situation where the ancient DNA community has been publishing two important bodes of data that cannot easily be co-analyzed by population genetic methods. To address this challenge, we introduce a subset of hundreds of thousands of SNPs for which 1240k data can be effectively co-analyzed with all other major data types.

---

The strategy of using artificially synthesized oligonucleotides as baits to fish out complementary sequences in a DNA library has been transformative in ancient human DNA studies. Enrichment has involved diverse approaches including oligonucleotides of various lengths affixed to glass slides (1), or baits that are free in solution (“in solution enrichment”) (2). Under appropriate chemical and temperature conditions, these baits hybridize to targeted molecules so that other molecules can be washed away, allowing the bound molecules to be isolated, released, and then sequenced. Enrichment has allowed researchers to achieve orders of magnitude enrichment for sequences that provide high information content to address important scientific questions.

In medical genetics, the most common application of target enrichment has been capturing the approximately two percent of the genome that constitutes the coding sequences of genes (the “exome”). When whole genome sequencing at high coverage was still prohibitively expensive, exome sequencing dropped the cost for surveillance of the coding regions for mutations causing rare diseases to affordable levels (2, 3). In ancient DNA analysis, the benefits of target enrichment are even greater (4). Not only is a tiny fraction of the genome in practice relevant for the great majority of analyses, but typically only a small proportion of molecules in the DNA library come from the individual of interest due to microbial contamination. Occasional ancient DNA libraries do contain most of their molecules from the individual whose bone or tooth was sampled, but it is more typical for most of the analyzed molecules not to be of human origin. For example, of more than 3,000 ancient individuals for which our research group published genome-wide data by the end of calendar year 2021, about half had less than 10% percent human DNA. Whole genome sequencing of such samples is prohibitively expensive for all but the most important samples given the typical budgets accessible to ancient DNA laboratories.

As an example of the challenge, consider a researcher who is interested in targeting a set of about 600,000 SNP positions genotyped in diverse modern human populations. Only perhaps one in a hundred ancient DNA sequences mapping to the human genome will overlap these positions, given the typical lengths of ancient molecules. If a DNA library is only one percent human, the proportion of sequences that will be informative for analysis will only be about one in ten thousand. Thus, if ~400 million DNA sequences are read from a library which is a typical number used to produce a ~30-fold whole human genome from modern DNA, only ~40,000 SNPs will be retrieved that overlap those genotyped in diverse modern populations. An individual with this much information will be only weakly informative for many analyses. In contrast, 25 million sequences from the same ancient DNA library after in-solution enrichment can provide coverage on nearly all targeted SNP positions by multiple unique molecules, allowing accurate inferences about population history at much lower cost.

Practical in-solution enrichment for ancient human DNA libraries was pioneered between 2010-2013 in studies that enriched for mitochondrial DNA (5, 6), nearly all of the unique sequences of chromosome 21 (5) and all or part of the exome (1, 7). In 2015, several papers were published that enriched for sequences overlapping sets of SNPs. The reagent that has been most extensively used and that we evaluate here is the ‘1240k reagent’: it targeted slightly fewer than 1.24 million SNPs chosen to be particularly valuable for studying variation among modern human populations (8–10). It has proven highly effective, and has been used to generate more than 70% of all genome-wide ancient human dataset: over five thousand individuals published in more than seventy papers (compiled at https://reich.hms.harvard.edu/allen-ancient-dna-resource-aadr-downloadable-genotypes-present-day-and-ancient-dna-data). The large body of data produced using the 1240k reagent has also created an important legacy dataset whose existence needs to be taken to account by researchers contemplating shifting to a new method: any future enrichment data benefits by targeting the same set of sites which can then be co-analyzed with existing data. However, the 1240k reagent also has limitations including variability in effectiveness of enrichment of targeted SNPs, and bias toward capturing some alleles more than others at the same sites, leading to technical artifacts when such data are co-analyzed with other data types such as random ‘shotgun’ sequencing data. Because of these technical challenges, researchers often restrict analyses to 1240k data only, or to shotgun data only, excluding key datapoints generated using the other strategy, and thus reducing the value of the world’s combined dataset.

A particular challenge with the 1240k reagent is that many ancient DNA laboratories have not been able to effectively access the technology. Secondary distribution of the reagent was not permitted by the company that synthesized the oligonucleotides. While the bait sequences were fully published in 2015, resynthesis of the reagent was prohibitively expensive on a per-reaction basis for laboratories interested in using the reagent on a scale of fewer than hundreds of samples. To make it possible for any ancient DNA researcher to carry out in-solution enrichment of more than a million SNPs, in 2021 two companies, Daicel Arbor and Twist Biosciences, made available in-solution enrichment reagents that target the core panel of 1.24 million SNPs as well as additional SNPs meant to address perceived gaps in the coverage of the original reagent. The co-authors of this study advised on the creation of these reagents, but were not paid as consultants and will not receive any remuneration from sale of the reagents. Here we describe a systematic comparison of all three reagents on a common set of 27 ancient DNA libraries chosen to span a range of library qualities from low to high percentages of human DNA, and from low to high complexities with respect to the number of unique human molecules present in the libraries (Table 1). In the interests of providing an independent assessment, our manuscript has not been reviewed by the companies that generated the reagents.

**Table 1:**
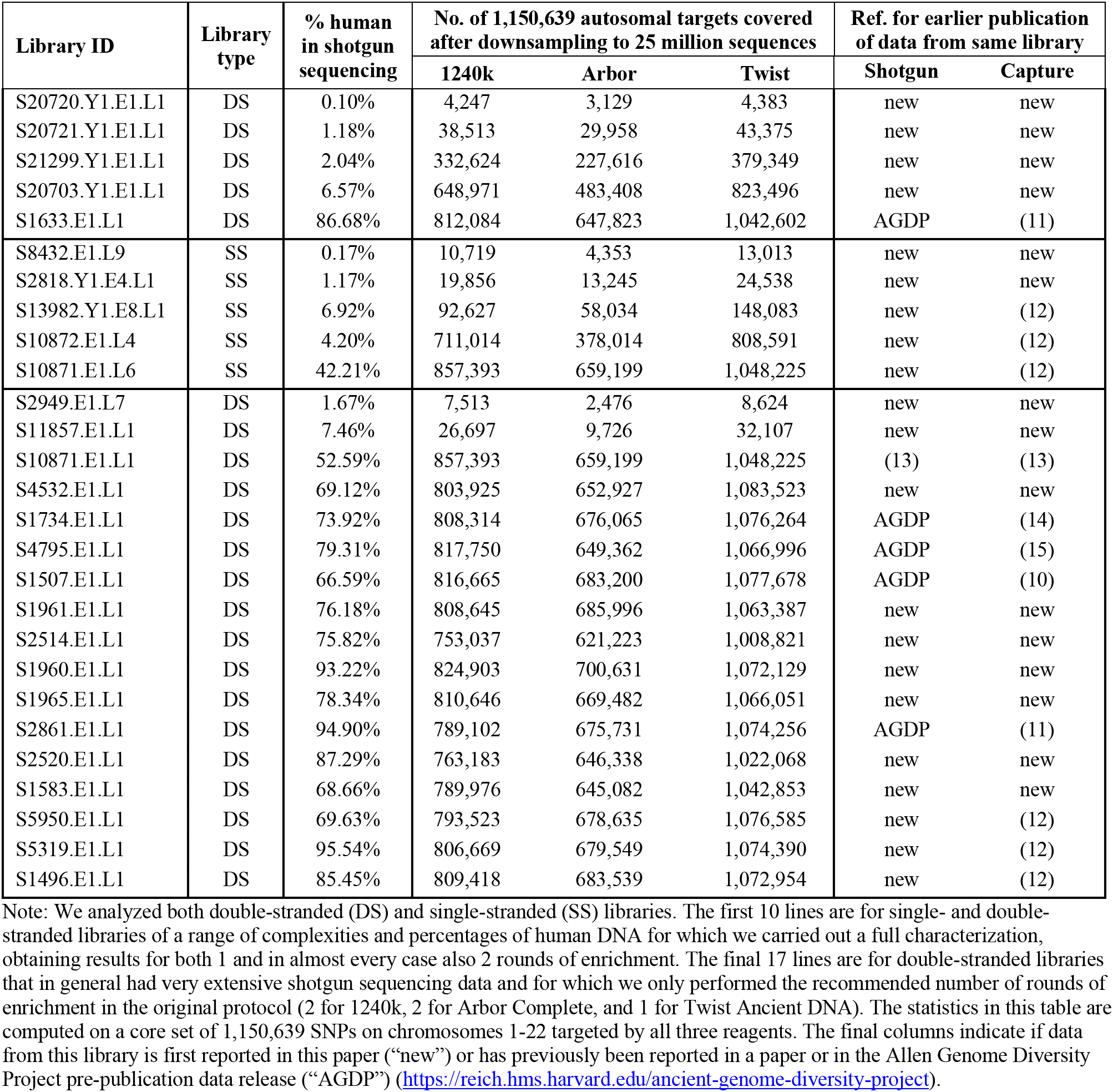
Twenty-seven ancient DNA libraries experimentally characterized in this study.

| Library ID | Library type | % human in shotgun sequencing | No. of 1,150,639 autosomal targets covered after downsampling to 25 million sequences |  |  | Ref. for earlier publication of data from same library |  |
| --- | --- | --- | --- | --- | --- | --- | --- |
|  |  |  | 1240k | Arbor | Twist | Shotgun | Capture |
| S20720.Y1.E1.L1 | DS | 0.10% | 4,247 | 3,129 | 4,383 | new | new |
| S20721.Y1.E1.L1 | DS | 1.18% | 38,513 | 29,958 | 43,375 | new | new |
| S21299.Y1.E1.L1 | DS | 2.04% | 332,624 | 227,616 | 379,349 | new | new |
| S20703.Y1.E1.L1 | DS | 6.57% | 648,971 | 483,408 | 823,496 | new | new |
| S1633.E1.L1 | DS | 86.68% | 812,084 | 647,823 | 1,042,602 | AGDP | (11) |
| S8432.E1.L9 | SS | 0.17% | 10,719 | 4,353 | 13,013 | new | new |
| S2818.Y1.E4.L1 | SS | 1.17% | 19,856 | 13,245 | 24,538 | new | new |
| S13982.Y1.E8.L1 | SS | 6.92% | 92,627 | 58,034 | 148,083 | new | (12) |
| S10872.E1.L4 | SS | 4.20% | 711,014 | 378,014 | 808,591 | new | (12) |
| S10871.E1.L6 | SS | 42.21% | 857,393 | 659,199 | 1,048,225 | new | (12) |
| S2949.E1.L7 | DS | 1.67% | 7,513 | 2,476 | 8,624 | new | new |
| S11857.E1.L1 | DS | 7.46% | 26,697 | 9,726 | 32,107 | new | new |
| S10871.E1.L1 | DS | 52.59% | 857,393 | 659,199 | 1,048,225 | (13) | (13) |
| S4532.E1.L1 | DS | 69.12% | 803,925 | 652,927 | 1,083,523 | new | new |
| S1734.E1.L1 | DS | 73.92% | 808,314 | 676,065 | 1,076,264 | AGDP | (14) |
| S4795.E1.L1 | DS | 79.31% | 817,750 | 649,362 | 1,066,996 | AGDP | (15) |
| S1507.E1.L1 | DS | 66.59% | 816,665 | 683,200 | 1,077,678 | AGDP | (10) |
| S1961.E1.L1 | DS | 76.18% | 808,645 | 685,996 | 1,063,387 | new | new |
| S2514.E1.L1 | DS | 75.82% | 753,037 | 621,223 | 1,008,821 | new | new |
| S1960.E1.L1 | DS | 93.22% | 824,903 | 700,631 | 1,072,129 | new | new |
| S1965.E1.L1 | DS | 78.34% | 810,646 | 669,482 | 1,066,051 | new | new |
| S2861.E1.L1 | DS | 94.90% | 789,102 | 675,731 | 1,074,256 | AGDP | (11) |
| S2520.E1.L1 | DS | 87.29% | 763,183 | 646,338 | 1,022,068 | new | new |
| S1583.E1.L1 | DS | 68.66% | 789,976 | 645,082 | 1,042,853 | new | new |
| S5950.E1.L1 | DS | 69.63% | 793,523 | 678,635 | 1,076,585 | new | (12) |
| S5319.E1.L1 | DS | 95.54% | 806,669 | 679,549 | 1,074,390 | new | (12) |
| S1496.E1.L1 | DS | 85.45% | 809,418 | 683,539 | 1,072,954 | new | (12) |

Note: We analyzed both double-stranded (DS) and single-stranded (SS) libraries. The first 10 lines are for single- and double-stranded libraries of a range of complexities and percentages of human DNA for which we carried out a full characterization, obtaining results for both 1 and in almost every case also 2 rounds of enrichment. The final 17 lines are for double-stranded libraries that in general had very extensive shotgun sequencing data and for which we only performed the recommended number of rounds of enrichment in the original protocol (2 for 1240k, 2 for Arbor Complete, and 1 for Twist Ancient DNA). The statistics in this table are computed on a core set of 1,150,639 SNPs on chromosomes 1-22 targeted by all three reagents. The final columns indicate if data from this library is first reported in this paper (“new”) or has previously been reported in a paper or in the Allen Genome Diversity Project pre-publication data release (“AGDP”) (https://reich.hms.harvard.edu/ancient-genome-diversity-project).

## Results

### Design of the three reagents

For completeness we begin by summarizing the original ‘1240k’ design, first reported in 2015 (8). The almost 1.24 million probes (1,233,013 after filtering to sites that could be robustly analyzed) were published in the supplementary materials of that study. Each SNP was targeted by four probes of 52 bp. To reduce bias toward capturing one allele or the other, two probes abutted but did not overlap the SNP in either direction. Another two probes were centered on the SNP, each with an alternative allele (again to reduce bias). The probes were appended on one side by an 8 bp universal flanking sequence and the 60 bp oligonucleotides were printed on Agilent 1M custom arrays. The baits were then biotinylated (5).

The SNP targets in the 1240k reagent were chosen to achieve a variety of purposes, summarized in Table 2 and the original publications (8–10). They included all the designable content of the Affymetrix Human Origins genotyping array (16) that has now been used to publish data on >8,800 present-day people from >840 human populations around the world (more than 90% of these data were published in thirteen studies (11, 16–28)). They included all the designable content of the Illumina 650Y genotyping array, part of a family of similar Illumina arrays whose content was iteratively optimized for genome-wide association studies and which have been widely used in genome-wide studies of human history. The 1240k targets furthermore included SNPs on the Affymetrix GeneChip Human Mapping 50K Xba Array; SNPs on the X chromosome to enable comparative studies of male and female population history; and SNPs on the Y chromosome to determine haplotypes. Finally, they included SNPs of phenotypic interest from association studies, scans of selection, or particularly important loci such as the HLA region of chromosome 6. In practice, 1240k reagent SNP enrichment experiments have also often include spiked-in oligonucleotide baits allowing enrichment of mitochondrial DNA (5, 6).

**Table 2:** Effectiveness of enrichment in different targeted subsets of the genome.

| Targeted subset of the genome<br>(some categories overlap) | # positions<br>(either SNPs or tiled<br>nucleotides) | 1240k<br>coverage<br>(vs. core set) | Twist<br>coverage<br>(vs. core set) | Arbor<br>coverage<br>(vs. core set) |
| --- | --- | --- | --- | --- |
| **SNPs** |  |  |  |  |
| Affymetrix Human Origins | 597,573 | 1.003 | 1.127 | 1.086 |
| Illumina 650Y | 660,611 | 0.951 | 0.882 | 0.927 |
| Affymetrix 50K | 58,559 | 0.371 | 0.516 | 0.71 |
| 1240k phenotypic supplement | 45,969 | 0.988 | 0.929 | 0.936 |
| 1240k X content | 49,704 | 1 | 1 | 1 |
| 1240k Y content | 32,670 | 1 | 1 | 1 |
| Twist phenotypic supplement | 94,587 | 0.059 | 0.943 | 0.136 |
| Twist Y content | 81,925 | 0.475 | 1.016 | 0.813 |
| Arbor ancestral supp. | 852,068 | 0.136 | 0.147 | 0.586 |
| Arbor Y supplement | 46,218 | 0.184 | 0.952 | 0.774 |
| **Tiling nucleotides** |  |  |  |  |
| Mitochondrial DNA | 16,569 | 6.17 | 2.955 | 28.51 |
| Twist HAR supplement | 857,339 (3171 HARs) | 0.039 | 2.448 | 0.09 |
| Twist gene sequencing supplement | 2,577 (in three genes) | 0.54 | 3.206 | 0.088 |
| Twist methylation targets | 80,000 (40,000 CpGs) | 0.086 | 3.584 | 0.109 |

Note: Relative coverage is computed by taking the average in this part of the genome after pooling data from all 27 libraries (2 rounds for 1240k, 2 rounds for Arbor, and 1 round for Twist), and dividing by either 1,150,639 SNPs on chromosomes 1-22, 49,704 SNPs on chromosome X (for SNPs there), or 32,670 SNPs for chromosome Y (for SNPs there). Coverage computations are based on sequence counts prior to removing PCR duplicated sequences.

For the Daicel Arbor “myBaits Expert Human Affinities” reagent, the oligonucleotide bait design is proprietary and the authors of this study do not have access to the technical details. Several modules are available (https://arborbiosci.com/genomics/targeted-sequencing/mybaits/mybaits-expert/mybaits-expert-human-affinities/). “Prime Plus” targets the exact same set of SNPs as the 1240k reagent along with the mitochondrial genome and a supplementary set of 46,218 Y chromosome SNPs. The “Complete” product adds an additional 852,068 transversion polymorphisms (“Ancestral Plus”) discovered as variable among archaic humans and validated as polymorphic in present-day humans (https://arborbiosci.com/wp-content/uploads/2021/03/Skoglund_Ancestral_850K_Panel_Design.pdf). These sites were chosen with the goal of facilitating analyses of African human population history, where biases due to the ancestry of the individuals in whom SNPs are discovered has the potential to complicate inferences (29). The fact that these SNPs are transversions is also useful when enriching ancient DNA libraries not enzymatically treated to remove ancient DNA damage which causes high error rates at transition SNPs. All the Arbor reagents also include baits to enrich mitochondrial DNA. We characterized the “Arbor Complete” reagent, which after accounting for the intersections of various SNP panels constitutes 2,131,299 SNPs.

For the Twist Biosciences “Twist Ancient DNA” reagent, a single 80 bp probe was centered on each targeted SNP. To avoid bias toward one allele or another, the nucleotide at the position of the SNP was chosen to be different from the two SNP alleles. The reagent was built around a core of 1,200,343 1240k SNPs (all 1240k SNPs on chromosomes 1-22 and X). It replaced the 32,670 1240k chromosome Y SNPs with 81,925 chosen to provide improved haplogroup resolution. It also added 94,586 additional phenotypically relevant targets chosen to target SNPs that were significant in genome-wide association studies in large sample sizes (32), as likely to have been affected by natural selection (33), as possibly implicated in rare disease (34), or as useful for computing heritability of complex traits (35) (Supplementary Section 1). These SNPs were only added to the reagent if they were not in high linkage disequilibrium with the core 1240k set (Supplementary Section 1, Online Table 1). The Twist reagent also targeted non-SNP locations. It tiled 857,339 bp in 3,171 Human Accelerated Regions (HARS); 2,577 bp in 3 genes relevant to α-thalassemia, β-thalassema, and favism; and 40,000 CpG dinucleotides where methylation rates are known to be correlated to human age (Supplementary Section 2). After filtering to probes that designed well, the final reagent included 1,434,155 probes targeting 1,352,535 SNPs. A mitochondrial panel from Twist Biosciences can be added to the bait pool; in practice we did not spike in sufficient concentrations of the mitochondrial DNA reagent to achieve consistently high mitochondrial DNA coverage, but subsequent experiments with more baits achieved results comparable to the other methods (data not shown).

### Empirical characterization of the three reagents

We experimentally characterized reagent performance in 27 libraries on which we performed 109 enrichment experiments (Table 1). All our sequencing was performed on HiSeqX10 instruments, and we report data on 12.2 billion merged sequences obtained for the enrichment experiments, and 43.3 billion merged sequences from shotgun sequencing. Basic statistics on the sequencing results for these libraries both before enrichment (shotgun sequencing), and after enrichment, are reported in Supplementary Table 1.

i. For 10 libraries of a range of complexities and percentages of endogenous human DNA (5 double-stranded and 5 single-stranded), we produced 0.006-26.7 mean coverage on the 1240k autosomal targets (assessed from 2 rounds of 1240k capture after removing duplicated molecules), and ranging in percentage of human DNA from 0.1% - 87%. We carried out 58 = 10 × 6 - 2 enrichment experiments on these libraries (the two most complex libraries were not captured for 2 rounds for Twist Ancient DNA). We carried out enrichment using all three reagents with the settings specified in the Methods, and deeply sequenced capture products both after the first and second round of sequencing, with 25-395 million merged sequences (median 95 million merged reads) for each experiment (Supplementary Table 1).
ii. For 17 double-stranded libraries 15 of which were of high complexity and high percentage of human DNA, we carried out extensive shogun sequencing, in 14 cases to more than 20-fold coverage. The shotgun data for four libraries has been fully published (12, 13, 36, 37), and the shotgun data for an additional 8 libraries has been released pre-publication as part of the Allen Ancient Genome Diversity Project / John Templeton Ancient DNA Atlas (https://reich.hms.harvard.edu/ancient-genome-diversity-project) (Table 1). We carried out 51=17×3 enrichments on these libraries with the experimental settings specified in the recommended protocols. Thus, we sequenced after two rounds of capture for 1240k and Arbor Complete, and one round of capture for Twist Ancient DNA. We sequenced the enriched products far more deeply than the ~25 million sequences typically generated for such experiments (median of 104 million merged sequences, Supplementary Table 1).

### Variation in effectiveness of enrichment in different parts of the genome

Table 2 highlights different targeted subsets of the genome, and shows the mean coverage in each category relative to the average achieved at the core set of 1,150,639 autosomal SNP positions (to assess coverage we use number of sequences obtained prior to removal of PCR duplicates as our goal here is to study the relative effectiveness of enrichment). In Online Table 1, we provide a SNP-by-SNP breakdown (this table also reports meta-information including why each SNP was targeted). Online Table 2 assesses the methylation targets (40,000 CpG dinucleotides). Online Table 3 covers Human Accelerated Regions (3,171 regions). Online Table 4 covers resequencing targets (in 3 regions). Online Table 5 reports 10.4 million alignable nucleotides on the Y chromosome. Online Table 6 reports results for 15,569 nucleotides of mitochondrial DNA.

All three methods not only enrich for the targeted content, but also for other positions usually within dozens of nucleotides on either side of explicitly targeted content (Figure 1). To obtain a better understanding of the patterns of enrichment near targeted locations and to assess if they can be useful, we annotated all 81.2 SNPs in the 1000 Genomes project dataset (38) by the coverage relative to the 1240k autosomal SNP targets (Online Table 7). All reagents effectively enriched not just the target SNPs, but hundreds of thousands of polymorphic positions nearby; for example, we identified ~130,000-170,000 SNPs that were enriched to ≥50% of the autosome-wide average coverage and had a minor allele frequency ≥5% in at least one 1000 Genomes Project continental population (Table 3). Researchers wishing to choose such non-targeted SNPs for inclusion in their analyses can select them based on the metrics in Online Table 7.

**Figure 1:**
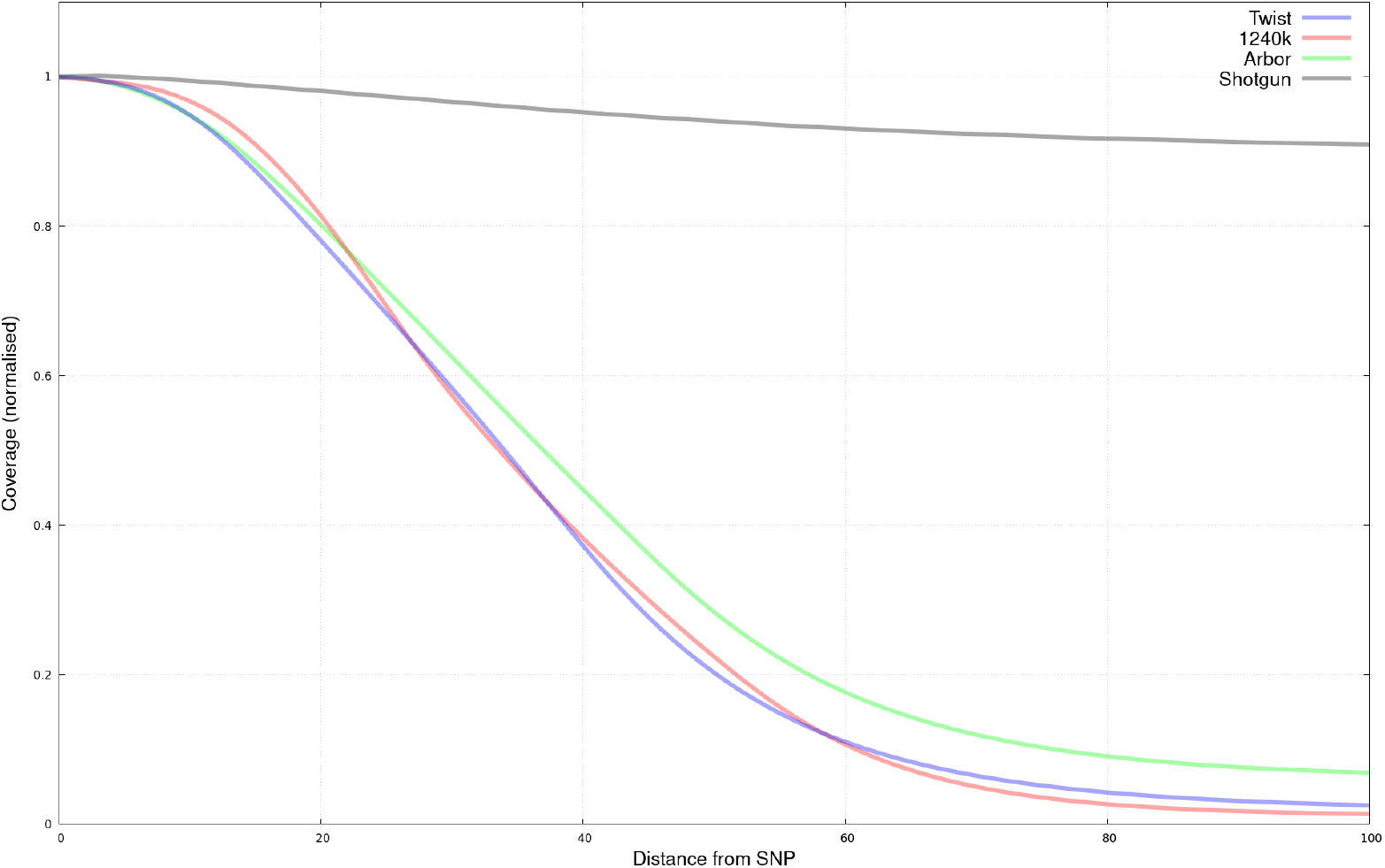
Distribution of sequence coverage as a function of distance from targets. Results are for the 15 high coverage sequencing libraries prior to removal of PCR duplicates, normalized by average coverage at targeted SNPs (position 0). Compared to nucleotides 100 base pairs from the closest target, coverage is 74-fold, 40-fold, and 15-fold enriched 1240k, Twist, and Arbor. Enrichment falls to 50% of targeted SNPs between 34-37 bases from SNP targets.

**Table 3:**
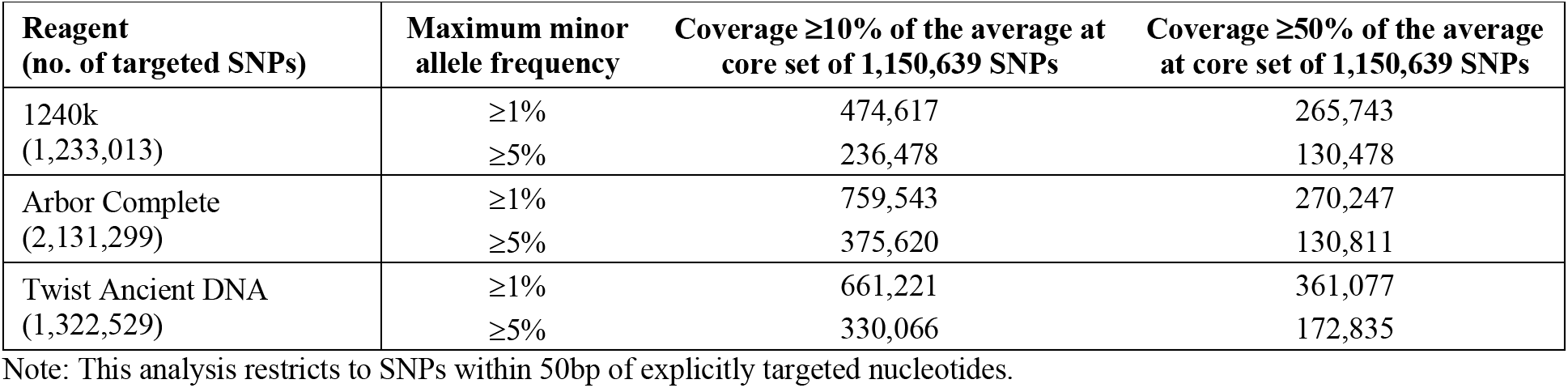
Enrichment of hundreds of thousands of near-target SNPs.

### Enrichment is less biased for Twist Ancient DNA than for other methods

We built histograms of coverage on targeted SNPs pooling over the libraries for which we had deep sequencing data (Figure 2A,B). The histograms are centrally peaked for shotgun sequencing (1% of SNPs with coverage <0.1-fold of the mean) and for Twist Ancient DNA (5% of SNPs), as expected for more homogeneous enrichment. In contrast, we observe skewed enrichment for 1240k (28% of SNPs with coverage <0.1-fold of the mean) and Arbor Complete (16% of SNPs).

**Figure 2:**
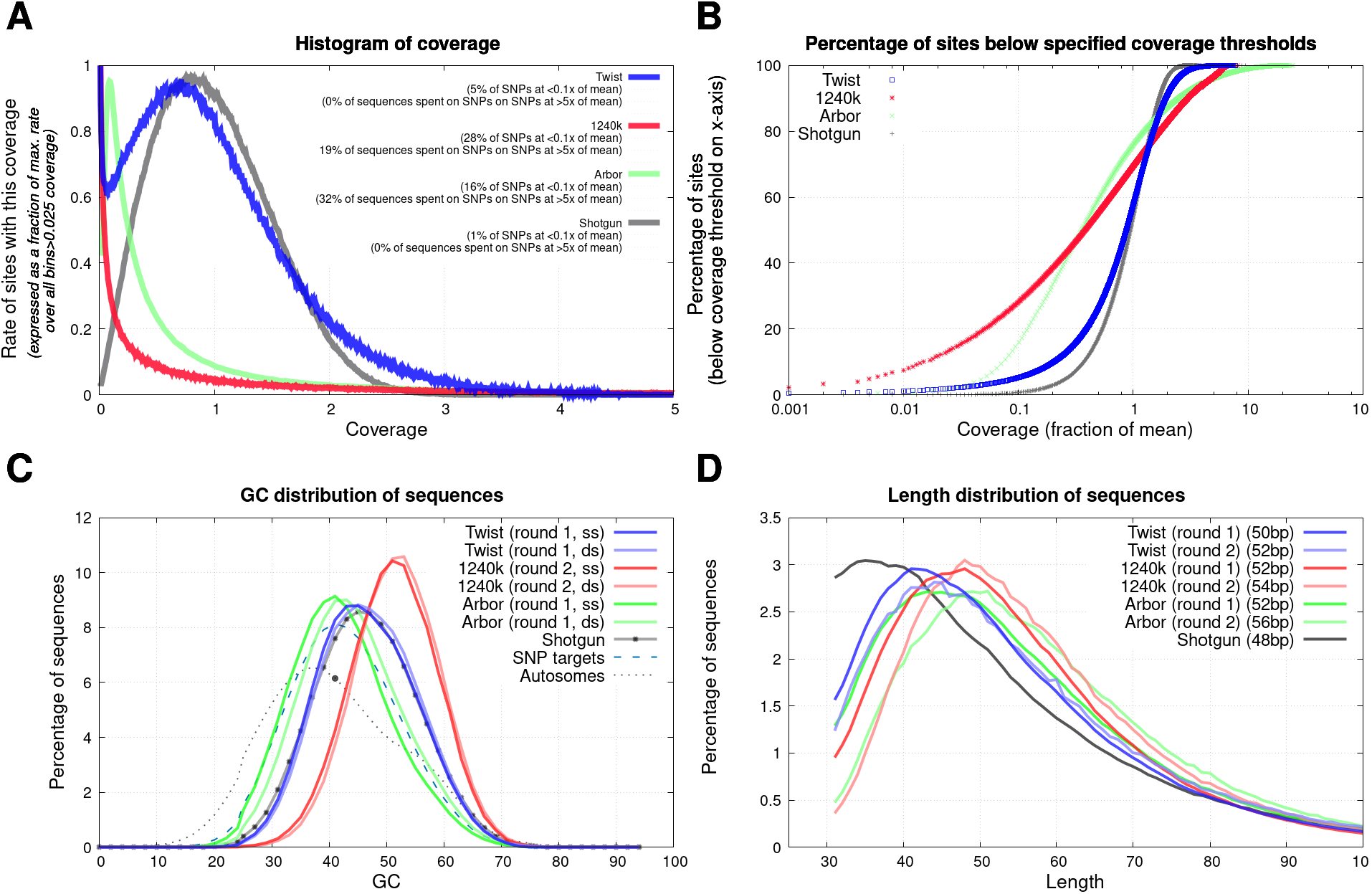
Biases in enrichment. We restrict to the 1,150,639 autosomal SNPs targeted by all three reagents. The top two panels analyze 15 libraries with high coverage shotgun sequencing data; the bottom two analyze 10 libraries with full results from both rounds of capture. (A) Variation in coverage across targeted SNPs is shown as a smoothed histogram where we normalize the y-axis by the maximum rate in bins with >0.025 of the average coverage. (B) The fraction of sites with coverage below different multiples of the mean. (C) The proportion of nucleotides that are either guanine or cytosine (GC) has a downward bias relative to the unenriched library for Arbor, an upward bias for 1240k, and little bias for Twist Ancient DNA. (D) All reagents preferentially enrich for longer molecules, with the least length effect for Twist Ancient DNA (medians for each data type are show in the legend).

Further evidence for a relatively homogeneous enrichment for Twist Ancient DNA comes from the proportion of guanines and cytosines in sequenced molecules, which is similar for Twist data and shotgun data, whereas Arbor Complete data shows a downward bias and 1240k an upward bias (Figure 1C). The Twist Ancient DNA also shows less of a bias toward an increase in the length of molecules than the other two enrichment methods (Figure 2D).

### All reagents are effective with Twist Ancient DNA consistently achieving highest coverage

As expected from its greater homogeneity in enrichment, Twist Ancient DNA achieves consistently high genome-wide coverage when measured by the number of SNPs hits at least once, for an amount of sequencing (25 million read pairs) that is typical for such experiments (Table 1). Compared to 1240k data the average increase in targeted SNP count is 1.21-fold, and compared to Arbor Complete it is 1.46-fold. We observe similar patterns for a range of sequencing coverages (Figure 3 and Supplementary Figure 1).

**Figure 3:**
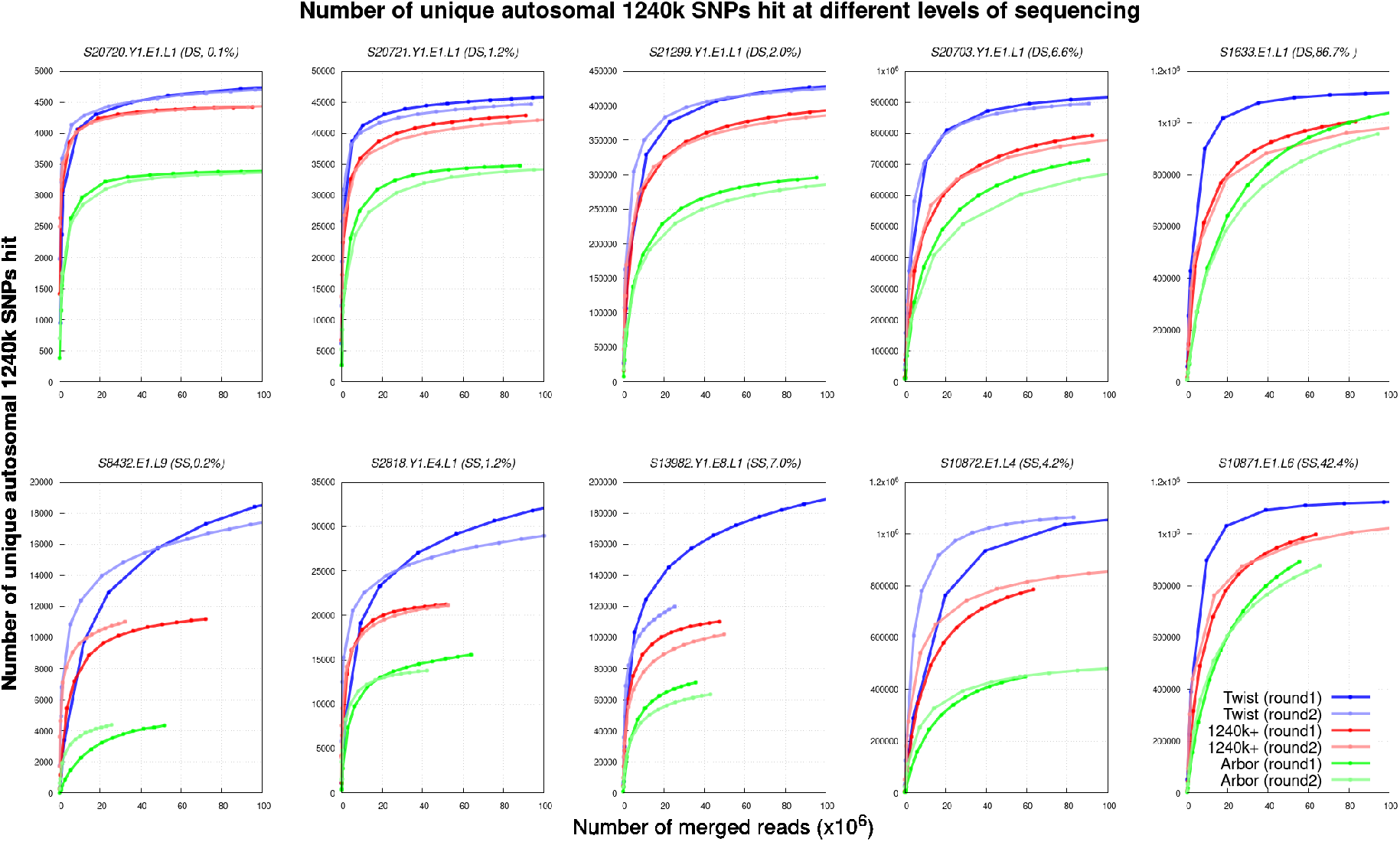
Performance of the three reagents over a range of sequencing depths. This analysis is based on various amounts of downsampling relative to the full sequencing data.

The increased yield for Twist Ancient DNA relative to the other protocols is particularly apparent for low complexity and single-stranded libraries, the condition for which we optimized this reagent over multiple rounds of testing. However, the Twist Ancient DNA reagent also outperforms the 1240k reagent for low-complexity double stranded libraries for which that methodology was optimized, highlighting how the Twist reagent is a definitively better reagent than 1240k from a technical point of view. For the Arbor Complete experimental settings we performed no optimization; instead, we used the manufacturer’s recommended protocol before product launch which differs from the one now available in the online manual. Better enrichment performance (perhaps much better) could likely be achieved with the Arbor Complete reagent through multiple rounds of optimization such as we performed for Twist Ancient DNA and 1240k. The correct lesson to take from these results is that the Arbor Complete reagent is effective and that these results place a minimum not a maximum on its utility.

A remarkable feature of all three enrichment method is the similar genome-wide coverage obtained from one round and two rounds of sequencing when a typical amount of data is collected after enrichment (~25 million sequences). This is striking in light of the fact that the proportion of sequences overlapping targets being much higher after two than one rounds of enrichment (average of 10-fold, median of 4-fold higher for the experiments in Figure 3) (Supplementary Table 1). The explanation is that the number of molecules typically sequenced after enrichment (~25 million), is far larger than the number of targeted positions. Thus, even with the relatively small proportions of molecules hitting targets after one round of enrichment, we in practice obtain sequences that cover the great majority of the targeted positions in the library. Because each enrichment round increases bias relative to the unenriched library, and because one round of enrichment is less expensive and time consuming than two, we recommend that standard practice for all three reagents should be to carry out just one round of enrichment (thus, the second round of enrichment for nearly all 1240k experiments to date was unnecessary).

A potential concern related to our approach of comparing results only at the 1,150,639 autosomal SNPs common to all three reagents is that this could be unfair to reagents that target more SNPs (especially Arbor Complete and to a lesser extent Twist Ancient DNA). In practice this is not a serious concern, as for a single round of enrichment which is our final recommended setting for all three reagents, the great majority of sequenced molecules miss targets (Supplementary Table 1), and thus the rate of molecules hitting targeted positions outside the 1,150,639 evaluation SNP set is small relative to the off-target content. In this setting, enrichment efficiency as assessed by the ratio of sequences overlapping the core set of SNP targets (the 1,150,639) to fully untargeted positions is similar to the same quantity if we do not drop sequences overlapping other targets. We use the number of merged sequences on the x-axis of Figure 3 instead of a corrected number, as number of merged sequences is intuitively understandable and relevant to real experiments.

### Addressing concerns about technical bias due to co-analysis of data from different sources

Biases associated with alignment and enrichment can affect population genetic analysis, causing data from two ancient DNA libraries processed using the same enrichment protocol to appear to have genetic affinities to each other even though the truth is that individuals from whom the libraries were obtained do not have distinctive relatedness. Concerns of this type have meant that in practice for population genetic analyses, researchers have often restricted their analyses to in-solution enrichment data using the 1240k reagent, or shotgun data, creating a challenging situation where two disjoint datasets have been built up in the community that are difficult to co-analyze. Even if a technology is more accessible to the community, and even if it is more efficient at capturing all targeted positions than the existing 1240k enrichment reagent, its practical value could be limited if it was difficult to co-analyze with data from other methods.

To explore how bias might affect our results, we began by projecting the data from the 15 libraries at the bottom of Table 1 onto a Principal Component Analysis of data from diverse present-day West Eurasian people living today (Figure 4). Encouragingly, all data from the same individuals plots at the same position, consistent with the pattern observed in the first publication of Twist Ancient DNA data where Neolithic individuals from Hazleton North in southern Britain clustered tightly whether the data source was 1240k or Twist (39). That study also showed that Twist and 1240k data could be robustly co-analyzed to detect familial relatedness (39).

**Figure 4:**
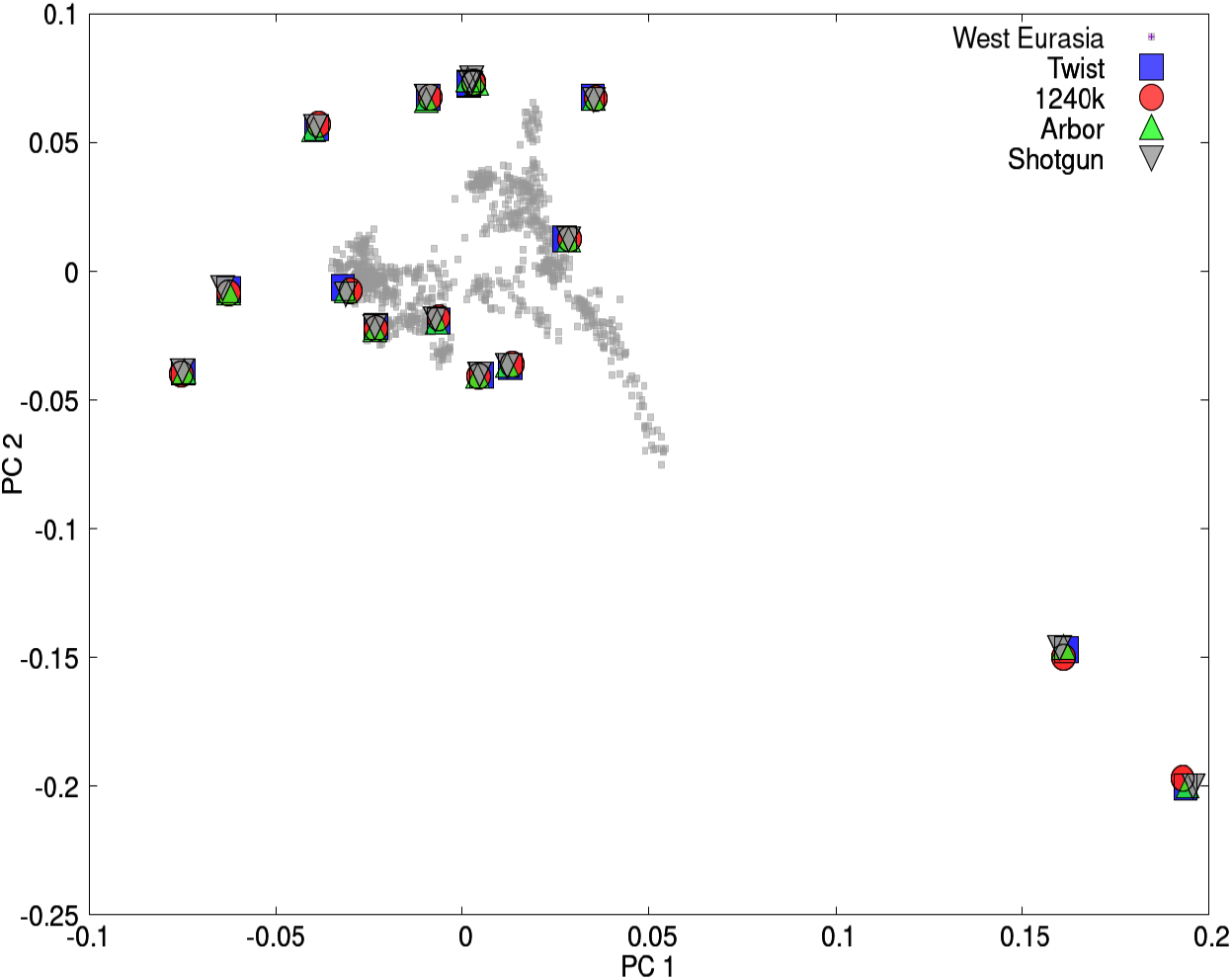
Principal Component Analysis shows similar ancestry regardless of data source. We performed PCA on >1000 West Eurasians, and projected data from the 15 individuals in the last rows of Table 1.

Lack of evidence for bias in PCA does not mean concerns about bias should be set aside. To further probe for bias associated with the different data generation technologies, for each of the 15 high coverage libraries we identified all SNP positions that were likely to be heterozygous based on observing at least one sequence matching both the reference allele and at least one matching the variant allele. For each SNP, we counted all reference and variant alleles observed at likely heterozygous positions beyond those not used in ascertainment; if there are no biases we expect 50% of these sequences to match the reference variant. We implemented an Expectation Maximization algorithm that uses these counts to estimate the distribution of reference bias for all SNPs, correcting for limited sample size which will produce more apparent variation in reference bias than is in fact the case (Supplementary Section 2).

We observe substantial average reference bias for all methods, which as expected due to the difficulty of mapping is word for shorter reads (Figure 5). A substantial degree of average reference bias is an important problem—and methods have been developed for mapping ancient DNA sequences in a way that reduces reference bias by an order of magnitude (40, 41)—but it is not the focus of this study, especially as reference bias also affects unenriched shotgun data. The unique issue for enrichment is the wider variation in reference bias across SNPs for 1240k and especially for Arbor Complete than for either shotgun or Twist Ancient DNA, even after controlling for sequence length (Figure 5). This reflects the fact 1240k and Arbor Complete, while not more likely to capture the reference allele on average, are more likely to skew from the mean degree of reference bias. Such skews specific to a technology are expected to cause data generated from two libraries processed by the same technology to have artifactual affinity.

**Figure 5:**
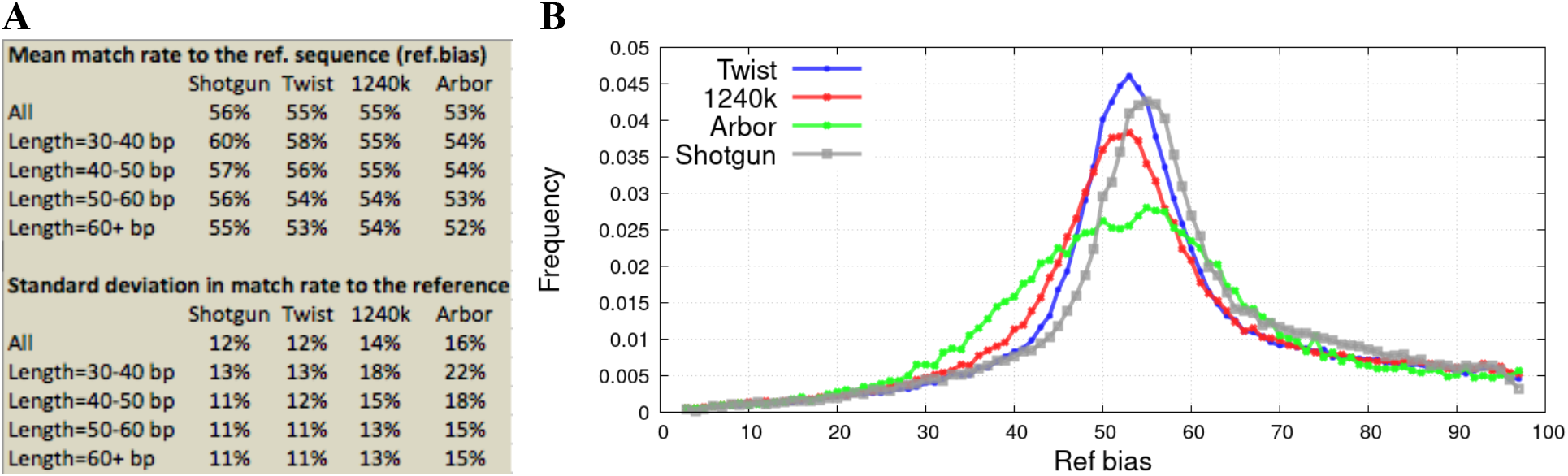
Allelic bias due to the different enrichment strategies. (A) Mean and standard deviation in the rate of matching to the reference genome for different data types, stratifying by sequence length, and correcting for stochastic error in the estimates using an Expectation Maximization (EM) algorithm described in Supplementary Section 2. (B) Distribution across SNPs in degree of reference bias. All analyses are based on sequences from loci ascertained as highly likely to be heterozygous, corrected for stochastic sampling variance using the EM.

To detect these artifactual attractions, we computed symmetry statistics of the form *f*_*4*_(library 1 - reagent A, library 1 - reagent B; library 2 - reagent A, library 2 - reagent B). If there are no technical biases, this quantity is expected to be 0, as data from each library should be symmetrically related to that from all other libraries. In contrast, if there are technical biases, we expect positive values of the statistic reflecting greater-than-random co-occurrences of alleles from two libraries processed using the same technology. Figure 6A computes a Z-score for the deviation of these *f*_*4*_-statistics from zero based on a Block Jackknife standard error; for the one-sided test appropriate here, Z>1.7 corresponds to P<0.05, and Z>3.1 corresponds to P<0.0001 (16). We observe that the Z-scores trend positive for all pairwise comparisons of the 15 libraries, as expected from the fact that any technical bias will cause a positive deviation. The statistics are most positive (mean Z of 3-4) for comparisons involving Arbor Complete captured SNPs, suggesting the strongest technical bias for this data type and consistent with the evidence that Arbor data has the largest standard deviation in reference bias across SNPs as shown in Figure 5A. The statistics are also large (mean Z almost 2) for statistics comparing 1240k to Twist Ancient DNA or shotgun data, as expected from the empirical observation of problems of co-analyzability of these two data types. Bias is minimal for Twist Ancient DNA comparisons to shotgun data (mean Z-score of around 0.6 with almost all Z-scores between −2 and 2) consistent with these two data types being far more co-analyzable from a population genetic perspective.

**Figure 6:**
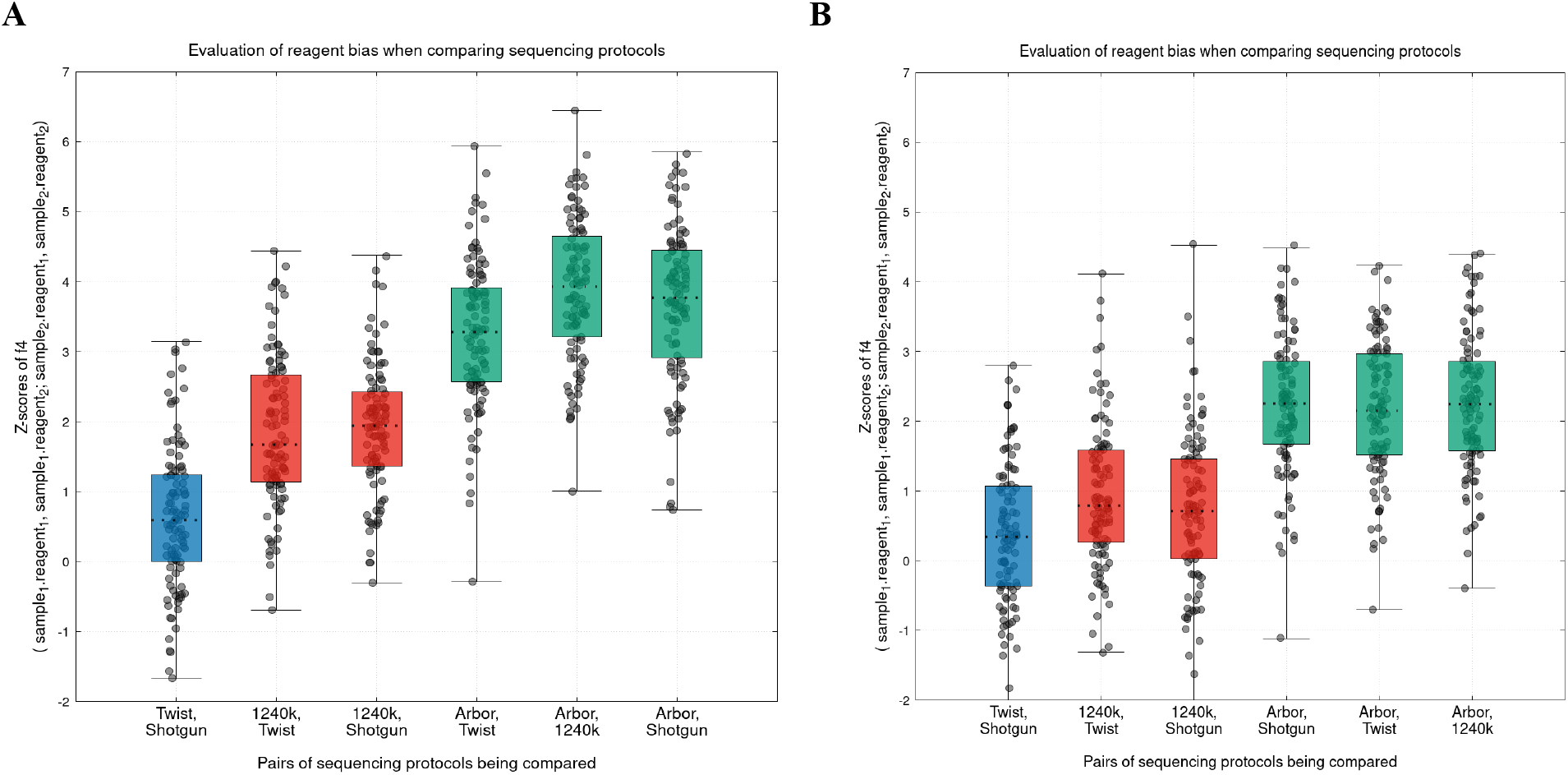
Artifactual attraction of data produced used the same methodology. (A) We compute symmetry statistics of the form *f*_*4*_(library 1 - reagent A, library 1 - reagent B; library 2 - reagent A, library 2 - reagent B), and plot Z-scores for all 105=15*14/2 pairwise comparisons of the 15 high coverage libraries as well as box-and-whisker plots showing full range, 25th and 75th percentiles, and mean. Statistics involving Arbor Complete are indicated with a green box; remaining comparisons involving 1240k with a red box; and the Twist Ancient DNA - shotgun comparison in blue. (B) Same analysis but restricted to a subset of 42% of autosomal SNPs chosen to have very similar rates of matching to the reference allele for shotgun and 1240k reagent data (empirically within 4% of each other).

While the minimal allelic bias associated with the data produced by the Twist Ancient DNA reagent and its easy co-analyzability with shotgun data addresses a limitation of the vast majority of capture experiments to date, it raises a new concern about co-analyzability of Twist Ancient DNA data with 1240k data. We therefore set out to identify a subset of SNPs with less susceptibility to such bias. To do this we mine data from 488 libraries for which we had shotgun data at a median of 5-fold coverage and also good 1240k data (much of this dataset is available as a pre-publication data release at https://reich.hms.harvard.edu/ancient-genome-diversity-project). We used imputation with GLIMPSE (42) to infer diploid genotypes at each SNP location (43) and counted rates of sequences matching to the reference and variant allele in all individuals where the posterior probability of being heterozygous was >0.9 at a given SNP. If there are no biases in enrichment, the frequency of observing the reference allele in the 1240k enrichment data is expected to match that in the shotgun sequence data (both 0.5). We restricted to the 42% of autosomal SNPs where difference in rate rates of matching to the reference allele for shotgun data and 1240k data was empirically less than 4% in the pooled reads over 488 libraries (this set of SNPs is specified as a column in Online Table 1). Figure 6B shows that the mean Z-scores for all *f*_*4*_-symmetry statistics comparing libraries that are shotgun sequenced, libraries enriched using 1240k, and libraries enriched using the Twist Ancient DNA reagent are between 0 and 1 after restricting to this set of SNPs, suggesting that this approach reduces biases.

Our goal in this analysis has been to demonstrate that a practical filter to reduce technical bias between methods exists; we have not attempted to optimize the filter and believe that there is substantial room to make the filter even better. The demonstration of the filter is also important for a reason that has nothing to do with Twist data, as it suggests a solution to problem that has been a long-standing challenge for ancient human DNA studies, namely, the difficulty of co-analyzing shotgun and 1240k enrichment data in genetic studies of population history. Applying a filter like has the potential to make data from diverse sources—1240k and shotgun and Twist— co-analyzable even for sensitive population genetic analyses.

## Discussion

We have systematically compared three in-solution reagents for enriching ancient DNA libraries for more than a million SNPs, and found that all three are highly effective.

The 1240k reagent has a proven track record, and has been used in more than 70 publications to report data from more than 5000 ancient individuals and to make robust inferences about population history. While 1240k data shows more allelic bias and less target homogeneiety than Twist Ancient DNA data, for studies of population history, the most important requirement is to regularly retrieve data from a large number of SNPs and it does this well.

The Arbor Complete reagent has several highly attractive features: it targets the same core set of SNPs as the 1240k enrichment reagent so that data can be co-analyzed, it targets an additional ~850,000 transversion SNPs chosen to be useful for studies of African population genetics, and it can be purchased commercially. Our implementation of Arbor Complete enrichment did not produce as high-quality results as the two other methods, but we also did not optimize the Arbor protocols in our lab as we did for the 1240k reagent and the Twist Ancient DNA reagent, and there is thus great potential for further performance improvement for this reagent.

The Twist Ancient DNA reagent was the most efficient of the three in our experiments, capturing sequences overlapping almost all targeted positions with relatively high homogeneity, achieving higher coverage, and having the least allelic bias making it most easily co-analyzable with shotgun data at nearly all analyzed SNPs. Like Arbor Complete, the Twist Ancient DNA reagent is commercially available. We have introduced a filter that makes it possible to tag SNPs most affected by the bias in 1240k enrichment, and which provides confidence that Twist data will be robustly co-analyzable with the great majority of ancient human DNA data generated to date.

Because of the multiple advantages associated with the Twist Ancient DNA reagent relative to 1240k in our testing, in June 2021 we performed our last of more than 28,500 1240k captures in our laboratory. Since then, we have performed more than 4,500 captures with Twist Ancient DNA reagent, and have already published our first data with this reagent (39). It is important for scientific communities periodically to update their methodologies when there are enough technical improvements, and we believe the advantages of new reagents are now so large that this time has come for ancient human DNA.

## Materials and Methods

### DNA extraction and library preparation

We extracted DNA from tooth or bone powder with a manual (44, 45) or automated protocol (46) using Dabney buffer and silica coated magnetic beads. We built the extract into indexed single-stranded USER-treated libraries (47) or into partial-UDG-treated barcoded double-stranded libraries (48). For cleanups after automated library preparation, we used silica coated magnetic beads and PB (Qiagen), and for cleanups after amplification we used SPRI.

### Target enrichment

The three target enrichment reagents all consist of biotinylated DNA probes, and while Arbor Complete and 1240k use single-stranded probes (52 bp for 1240k, unknown to us for Arbor Complete), Twist Ancient DNA uses double-stranded 80 bp probes. The original protocol for Twist reagents specified one round of enrichment, whereas the original protocols for Arbor Complete and 1240k specified two consecutive rounds of enrichment. Arbor Complete and 1240k had the mitochondrial panel included in our testing (1240k reagent: 3 bp tiled probes of mitochondrial genome of 52 bp length, spiked in at 0.033%), whereas for Twist Ancient DNA we only added the Twist Mitochondrial Panel to 19 of the 27 libraries (120 bp long probes, spiked in at 1.67%). In our Twist testing, we added in the mitochondrial DNA probes at a tenth of the concentration we had intended (our plan had been to spike in at 16.7% but effectively we used 10x less because the concentration in the kit was 10x lower than expected). In subsequent experiments with the intended concentration, we have obtained more efficient mitochondrial retrieval for Twist than we show in Online Table 6.

For a total of 10 ancient human DNA libraries (5 single-stranded and 5 double-stranded) of varying genomic complexity and endogenous content (Table 1), we enriched for one and in almost every case two rounds following the conditions below for each enrichment reagent. Additionally, we enriched 15 high-complexity libraries and 2 low-complexity libraries for which we had generated large amounts of shotgun sequence data to further investigate the performance of each reagent. For these libraries, we used only the originally recommended settings: 1 round for Twist Ancient DNA, 2 rounds for 1240k, and 2 rounds for Arbor Complete.

### 1240k reagent

Since the development (5) of the in-solution enrichment technology that is the basis for the 1240k reagent, we have changed temperature settings in our implementation, but not buffer composition or volumes. For this study, we started with 1 μg of library and hybridized to 1 μg of single-stranded biotinylated bait in a total volume of 34 μl for at least 16 h at 73 ℃. We bound the biotinylated probes to 30 μl MyOne streptavidin C1 beads in Binding Buffer for 30 min, and washed the beads 5 times with 3 different wash buffers (stringent washes were performed 3 times at 57 ℃). We melted the library molecules from the probes, precipitated onto magnetic beads, washed, eluted and amplified for 30 cycles using appropriate primer pairs (depending on whether they were single- or double-stranded libraries) and Herculase II Fusion polymerase. We cleaned up the product with 38% SPRI reagent and eluted round 1 in 15 μl TE. For round 2, we used 5 μl of the round 1 product (usually 500-700 ng total) and hybridized with 500 ng of single-stranded baits again for about 16 h. Capture and washes were identical to round 1, but we eluted the cleaned PCR product in 50 μl usually resulting in 50 - 90 ng/μl product.

### Arbor Complete

We used the ‘myBaits Expert Human Affinities - Complete panel’. The kit was not commercially available at the time of testing, and we therefore used reagents and buffers also used for 1240k as recommended by representatives of Daicel Arbor. Experimental settings are similar to the 1240k settings, with the following adjustments. Hybridization was performed at 70 ℃ and binding to 30 μl MyOne Streptavidin beads in binding buffer was recommended at 70 ℃ for 5 min. All washes were identical to 1240k, but the 3 stringent washes were performed at 55 ℃ and amplification cycles were reduced to 20 in round 1. The entire product was used in round 2 (except for the 10 libraries we tested 1 and 2 rounds of capture, 1/7th was kept for round 1 indexing PCR and sequencing) and the final amplification was only performed for 12 cycles. The now commercially available kit is slightly different and the recommended settings can be found online (https://arborbiosci.com/wp-content/uploads/2021/03/myBaits_Expert_HumanAffinities_v1.0_Manual.pdf).

### Twist Ancient DNA

We explored a range of probe lengths, reagent volumes and temperature settings to optimize performance for unmultiplexed low-complexity single stranded ancient DNA libraries. The experimental conditions which used here (which are substantially different from the protocol optimized by Twist for in-solution enrichment products applied to multiplex modern DNA) are as follows. We used 1 g of dried library and reconstituted in 7 μl of Universal Blockers and 5 μl Blocker Solution. In a second plate, we combined 5 μl of Hybridization mix (standard protocol is 20 μl) with 1 μl of Twist Ancient DNA probes (this is an optimized volume based on our testing; the standard protocol from Twist for modern high quality DNA specifies 4 μl). We melted the (double-stranded) probes for 5 min at 95 ℃ and cooled to 4 ℃ for 5 min. During the 4 ℃ cooling of the probes, we incubated libraries and blockers for 5 min at 95 ℃. We next equilibrated both plates to room temperature for 5 min. We added the 6 μl of probe (6.167 μl if mitochondrial DNA probes were added) and hybridization buffer to the 12 μl library and blocker, mixed, and overlaid with 30 μl Hybridization Enhancer and incubated at 62 ℃ (standard is 70 ℃) in a thermal cycler for at least 16 h. We used 300 μl Streptavidin beads (standard is 100 μl) and bound for 30 min at room temperature. In manual processing, we next washed beads 4 times with 2 different wash buffers; of these, 3 were stringent washes at 49 ℃ (standard is 48 ℃) (in automated processing, we performed 7 washes of which 6 were stringent washes at 49 ℃; the automation protocol is available from Twist Biosciences). We amplified from 50% of the bead slurry with Kapa HiFi HotStart ReadyMix for 23 cycles (standard is fewer cycles) with the provided primers (ILMN) for single-stranded libraries or indexing primer for double-stranded libraries in an off-bead PCR. We finished by purifying the PCRs with 1.8x Purification Beads (standard is 1x) and eluted in 50 μl TE.

### Sequencing

We sequenced enriched and shotgun libraries on HiSeqX10 instruments with 2×101 cycles, and either 2×7 cycles (double-stranded libraries) or 2×8 cycles (single-stranded libraries) to read the index sequences.

### Bioinformatic data processing

Because the enriched ancient DNA libraries were sequenced in pools, we needed to demultiplexed sequences which we did based on two different types of oligonucleotide tags: library-specific barcode pairs (for double-stranded libraries) and index pairs (for all libraries). We merged paired-end sequences requiring either a minimum of 15 base pair overlap (with at most one mismatch, base quality≥20) or up to three mismatches of lower base quality. We mapped these sequences to the human genome (*hg19)* using *samse* from *bwa-v0.6.1* (49). We restricted analysis to merged sequences of at least 30 base pairs. For analyses in which we were interested in relative efficiency of retrieval of molecules at different targeted locations, we measured coverage prior to removal of PCR duplicated molecules; for other analyses, we assessed coverage after removal of PCR duplicates. To represent each nucleotide position for analyses that required SNP genotype calls (Principal Component Analysis and *f*_*4*_-statistics), we chose a random sequence at each location, requiring a mapping and base quality of 10 and 20.

### Fraction of published ancient DNA data produced by in-solution enrichment

To compute the proportion of genome-wide ancient human DNA data for which data had been generated by 1240k enrichment (>70%), we used all published data from version v51 of the Allen Ancient DNA Resource (https://reich.hms.harvard.edu/allen-ancient-dna-resource-aadr-downloadable-genotypes-present-day-and-ancient-dna-data), consisting of compiled records of published genome-wide ancient human DNA data as of December 22, 2021.

### Distribution of endogenous DNA proportion in published ancient DNA data

To compute the fraction of individuals with proportions of endogenous DNA below different thresholds, we restricted to published data from our laboratory for which we had at least 15,000 SNPs on chromosomes 1-22 present targeted by the 1240k reagent, and assessed as passing quality control either fully (‘PASS’) or with minor concerns (‘QUESTIONABLE’). We restricted to individuals for which we had an endogenous DNA proportion estimate for at least one library, and represented each individual by the library with the highest proportion of endogenous DNA.

## Data Availability Statement

The aligned sequences are available through the European Nucleotide Archive, accession [to be made available upon publication].

## ACKNOWLEDGMENTS

We thank Kim Callan, Elizabeth Curtis Lora Iliev, Lijun Qiu, Noah Workman, and Fatma Zalzala for support in the wet laboratory. We are grateful to Mark Consugar, Ellie Juarez, Paul Frere, Keith McKenna and Frank Capriglione at Twist Biosciences who supported the development of the Twist Ancient DNA reagent. We thank Ryan Doan, Steve Horvath, Iosif Lazaridis, Alissa Mittnik, Vagheesh Narasimhan, and Iñigo Olalde, who advised on choice of additional SNPs and targeted regions for the Twist reagent, and Ali Akbari who created the imputed dataset that made it possible to identify SNPs with reduced susceptibility to capture bias. We thank Jacob Enk and Alison Default at Daicel Arbor who drove the development of the myBaits Expert Human Affinities capture reagents; and Pontus Skoglund and Yassine Souilme who advised on SNP choice for that reagent (none of these colleagues had input into the manuscript). We thank Songül Alpaslan-Roodenberg, Ian Armit, Nihat Erdogan, Julian Jansen van Rensburg, Carles Lalueza-Fox, Benjamin Neil, Ron Pinhasi, Mary Prendergast, Bob Sattler and Irina Shingiray for the collaborations that produced the ancient DNA data samples used for the technical comparisons reported in this study; this paper does not provide information on archaeological context of the analyzed libraries, although such analyses were previously reported for some individuals (Table 1). This research was funded by NIH grants GM100233 and HG012287, by the Allen Discovery Center program, a Paul G. Allen Frontiers Group advised program of the Paul G. Allen Family Foundation, by John Templeton Foundation grant 61220, and by the Howard Hughes Medical Institute.

## Supplementary Information Summary

### Supplementary Tables

Supplementary Table 1 Sequencing results on all 27 libraries

### Supplementary Figures

Supplementary Figure 1 10 library downsampling experiment using coverage as output

### Supplementary Information

Supp. Information section 1 Content added to Twist Ancient DNA Reagent beyond 1240k

Supp. Information section 2 EM Algorithm to Correct for Binomial Sampling Variance

### Online Tables (large text files, all compressed)

Can be accessed through the following Dropbox link: https://www.dropbox.com/sh/h024odwt5w1yc37/AAC9jCMhhOncXQRBaWMWOzPla?dl=0

|  |  |  |  |
| --- | --- | --- | --- |
| Online Table 1 | Twist SNP targets | 1,352,529 rows | SNPs |
| Online Table 2 | Methylation CpG targets | 80,000 rows | bases |
| Online Table 3 | Human Accelerated Regions (HAR) | 857,339 rows | bases |
| Online Table 4 | Gene Resequencing Regions | 2,577 rows | bases |
| Online Table 5 | Mappable Y Chromosome | 10,446,037 rows | bases |
| Online Table 6 | Mitochondrial DNA | 16,569 rows | bases |
| Online Table 7 | Statistics at 1000 Genomes SNPs | 81,286,436 rows | SNPs |

**Supplementary Table 1:**
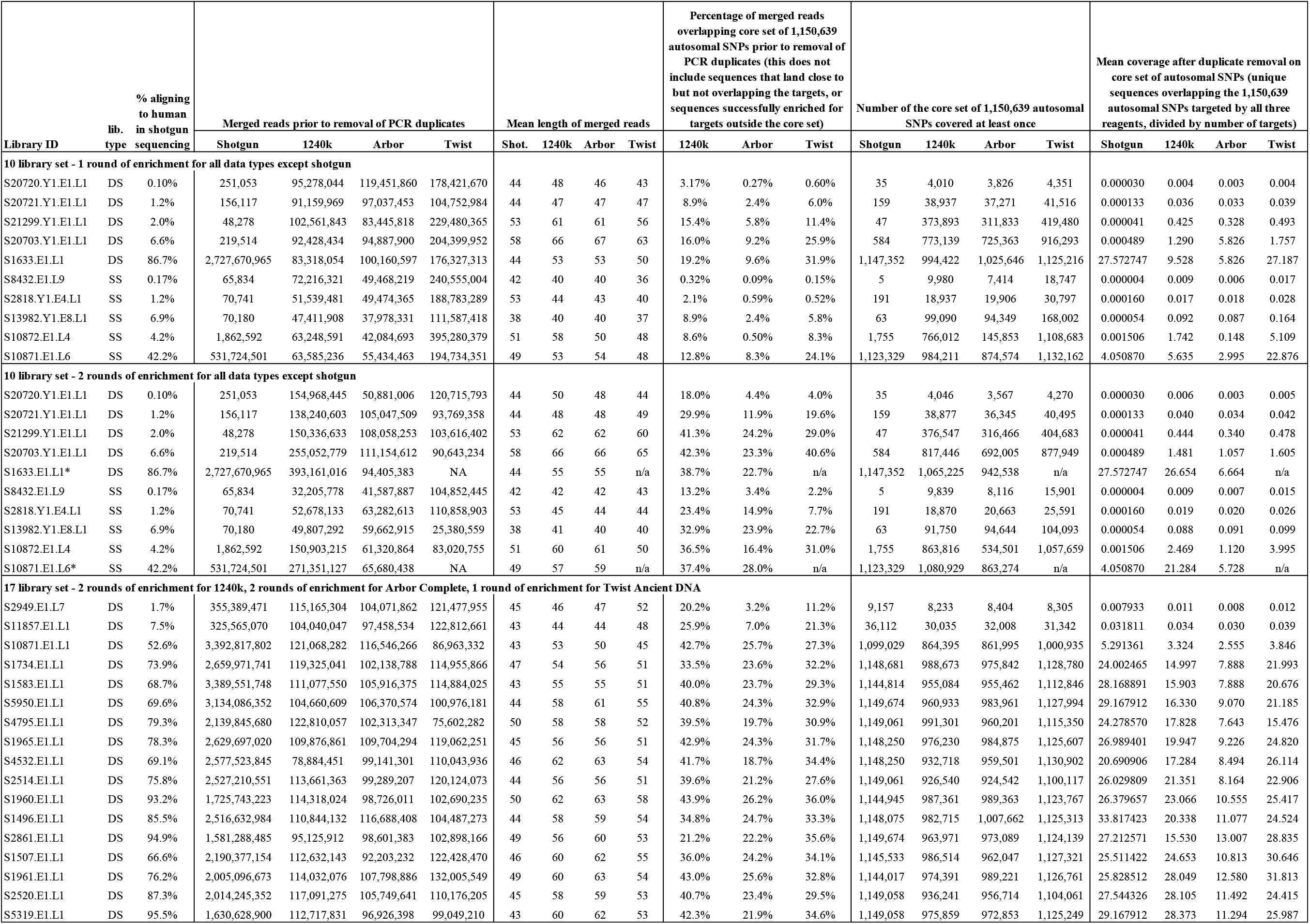
Sequencing results on all 27 libraries. A total of 10 libraries were sequenced after both the first and second round of enrichment (except for S1633.E1.L1 and S10871.E1.L6 which were not sequenced after a second Twist round). The bottom 17 libraries reflect 2, 2 and 1 rounds of enrichment for 1240k, Arbor and Twist respectively. DS - double-stranded, SS - single-stranded.

**Supplementary Figure 1:**
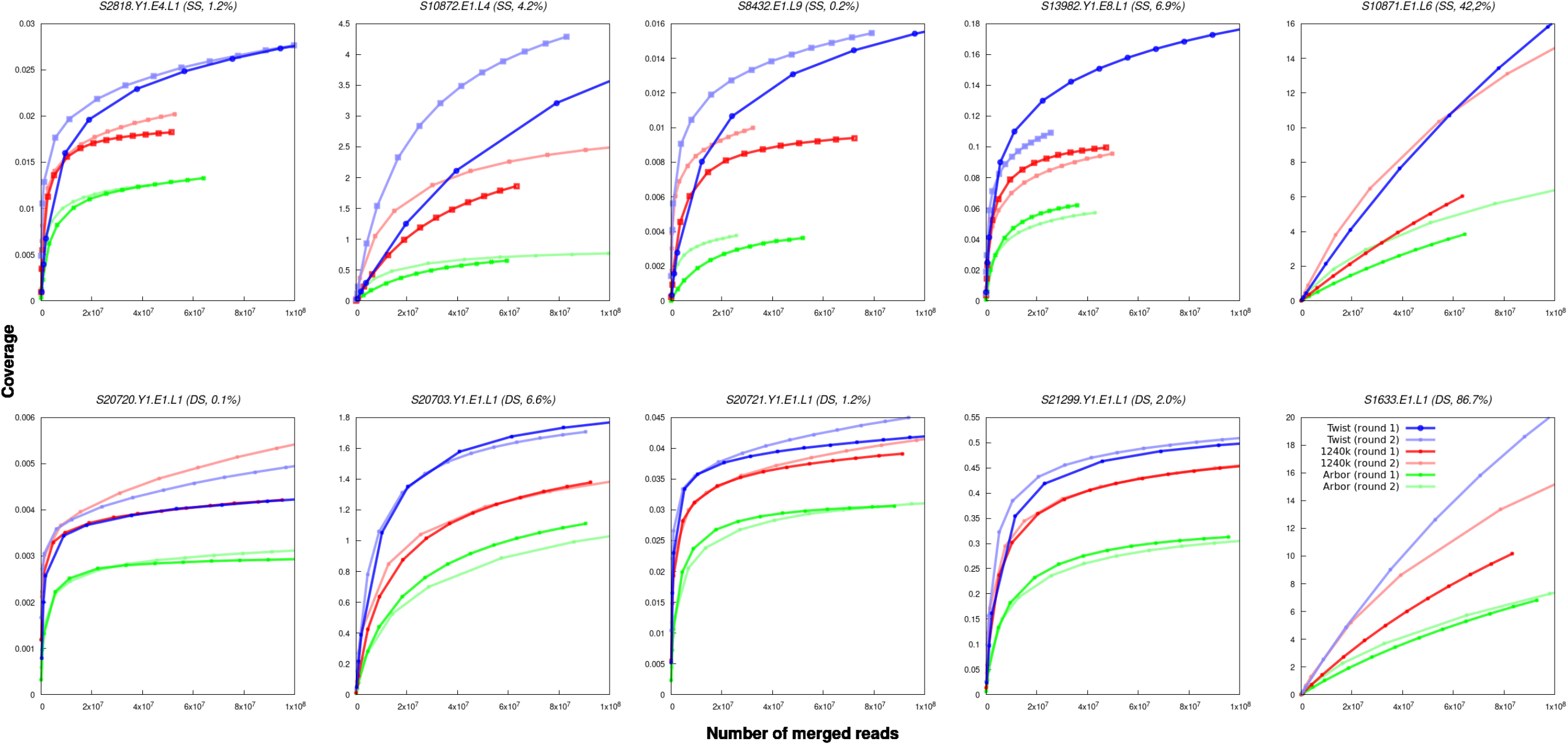
10 library downsampling experiment using coverage as output.

### Supplementary Section 1: Content added to Twist Ancient DNA Reagent beyond 1240k

#### (1a) Adding 94,586 polymorphisms on chromosomes 1-22 and X

For the Twist Ancient DNA reagent, we began by attempting to bait all 1,233,013 SNPs in the 1240k reagent. We then added additional content to target SNPs of phenotypic significance or SNPs improving characterization of variation on the Y chromosome.

- *“GWAS” SNPs (SNPs associated with phenotypes in Genome-Wide Association Studies)* We used a list of 236,638 SNPs that are genome-wide significant in one of 4,155 GWAS’s on 558 traits in a diverse set of populations (32). In contrast to the GWAS catalog database (50), this list only includes SNPs identified in GWAS of 50,000 individuals or more.
- *“RELATE” SNPs* We included SNPs estimated to have been under recent selection in any of 26 diverse modern populations from the 1000 Genomes Project (38) based on distortions in coalescent tree shapes (33). We selected all 61,308 SNPs with selection p-values < 10^−5^ in any population.
- *“Clinvar” SNPs* We included 32,689 SNPs from the Clinvar database by selecting all variants where the highest reported allele frequency is >1% (34) (https://www.ncbi.nlm.nih.gov/clinvar/). These SNPs are highly enriched for coding, non-synonymous variants.
- *“Polyfun” SNPs* We included 75,592 fine-mapped SNPs falling in regions with functional annotations that are enriched for heritability for a range of complex traits, specifically all SNPs with Posterior Inclusion Probability of >0.1 (35).

#### (b) Linkage disequilibrium (LD) pruning to remove genetically correlated SNPs

We pruned the selected SNPs for linkage disequilibrium in 2,261 individuals from the 1000 Genomes Project. For pruning, we use the PLINK (51) command --indep-pairwise 1000 100 0.9.

We computed LD for each of the remaining SNPs to the core set of 1240k SNPs using the command -- r2 --ld-window-r2 0.2 --ld-window 10 --ld-window-kb 1000. We excluded all SNPs with LD greater than 0.9 to any 1240k SNP.

#### (c) Quality control

We characterized SNPs from all sources by their dbSNP reference numbers (rs-IDs) as well as their reference and variant alleles. We filtered out insertion/deletion polymorphisms. We mapped rs-IDs to chromosome and position and determined alleles using the Ensembl database for genome build GRCh37 (hg19), accessed through biomaRt (http://www.biomart.org/). The hg19 reference sequence (“hg19_1000g.fa.gz”) was then used to obtain 52 bp flanking either side. For multi-allelic sites, the two variants identified in the original sources were kept. Alleles in the hg19 reference sequence were designated as “ref”, and the alternative alleles as “alt”.

Table S1.1 shows a record of the SNPs deriving from each of these four methodologies, including the number retained after the different pruning steps; this identified 94,586 SNPs.

**Table S1.1:** SNPs selected from each source (there is some overlap, so total is not the sum)

| Name | Initial | Not in 1240k | After pruning | $R^2 < 0.9$ | Would keep | Mean allele frequency | Mean $R^2$ ($> 0.2$) | Mean $R^2$ ($\leq 0.2$) |
| --- | --- | --- | --- | --- | --- | --- | --- | --- |
| Clinvar | 32705 | 27495 | 20544 | 17262 | 17601 | 0.167 | 0.7 | 0.337 |
| GWAS | 236638 | 160819 | 66857 | 38540 | 38478 | 0.401 | 0.79 | 0.012 |
| Polyfun | 75592 | 59500 | 42088 | 32430 | 33145 | 0.279 | 0.72 | 0.174 |
| Relate | 61308 | 49701 | 23228 | 14579 | 14428 | 0.419 | 0.78 | 0.008 |
| Total | 375408 | 276824 | 140520 | 93812 | 94586 | 0.361 | 0.77 | 0.066 |

Note: "Would keep" includes SNPs not in the 1000 Genomes Project and with unclear LD, and excludes SNPs with mismatching alleles or positions.

We sought to understand the genomic distribution and other characteristics of the newly added SNPs. Table S1.2 shows the distribution across chromosomes for each of the four methodologies. Figure S1.1 shows the allele frequency distribution of the variant allele. Figure S1.2 shows the distribution of maximum R^2^ to any 1000 Genomes Project SNPs.

**Table S2:** Number of newly targeted SNPs by chromosome.

| Chromosome | Clinvar | GWAS | Polyfun | Relate |
| --- | --- | --- | --- | --- |
| 1 | 1496 | 2932 | 3053 | 1052 |
| 2 | 1612 | 4291 | 2773 | 1348 |
| 3 | 890 | 2775 | 2094 | 989 |
| 4 | 693 | 1902 | 1497 | 922 |
| 5 | 922 | 2486 | 1826 | 830 |
| 6 | 908 | 3020 | 1903 | 764 |
| 7 | 741 | 2030 | 1883 | 800 |
| 8 | 668 | 2300 | 1279 | 820 |
| 9 | 907 | 2158 | 1391 | 907 |
| 10 | 684 | 1549 | 1479 | 702 |
| 11 | 1031 | 2066 | 1677 | 714 |
| 12 | 888 | 2019 | 1841 | 644 |
| 13 | 392 | 999 | 909 | 458 |
| 14 | 533 | 1036 | 910 | 527 |
| 15 | 613 | 1517 | 1149 | 476 |
| 16 | 1042 | 1081 | 1433 | 713 |
| 17 | 1095 | 1156 | 1646 | 380 |
| 18 | 341 | 785 | 771 | 379 |
| 19 | 936 | 607 | 1557 | 307 |
| 20 | 464 | 1204 | 1098 | 301 |
| 21 | 324 | 80 | 410 | 192 |
| 22 | 378 | 485 | 567 | 203 |
| X | 43 | NA | NA | NA |
| Total | 17601 | 38478 | 33146 | 14428 |

**Figure S1.1:**
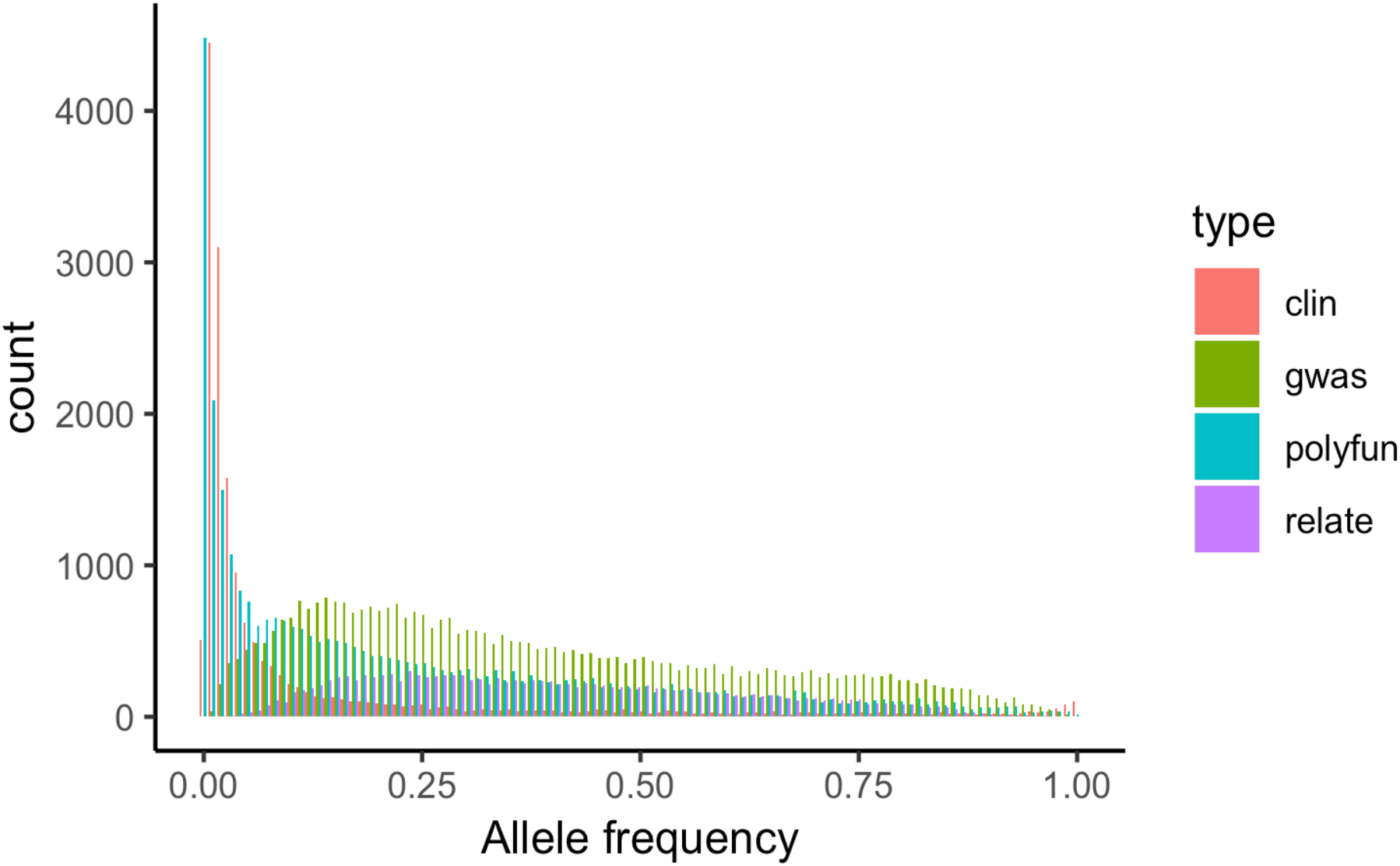
Allele frequency distribution by source of newly added SNPs.

**Figure S1.2:**
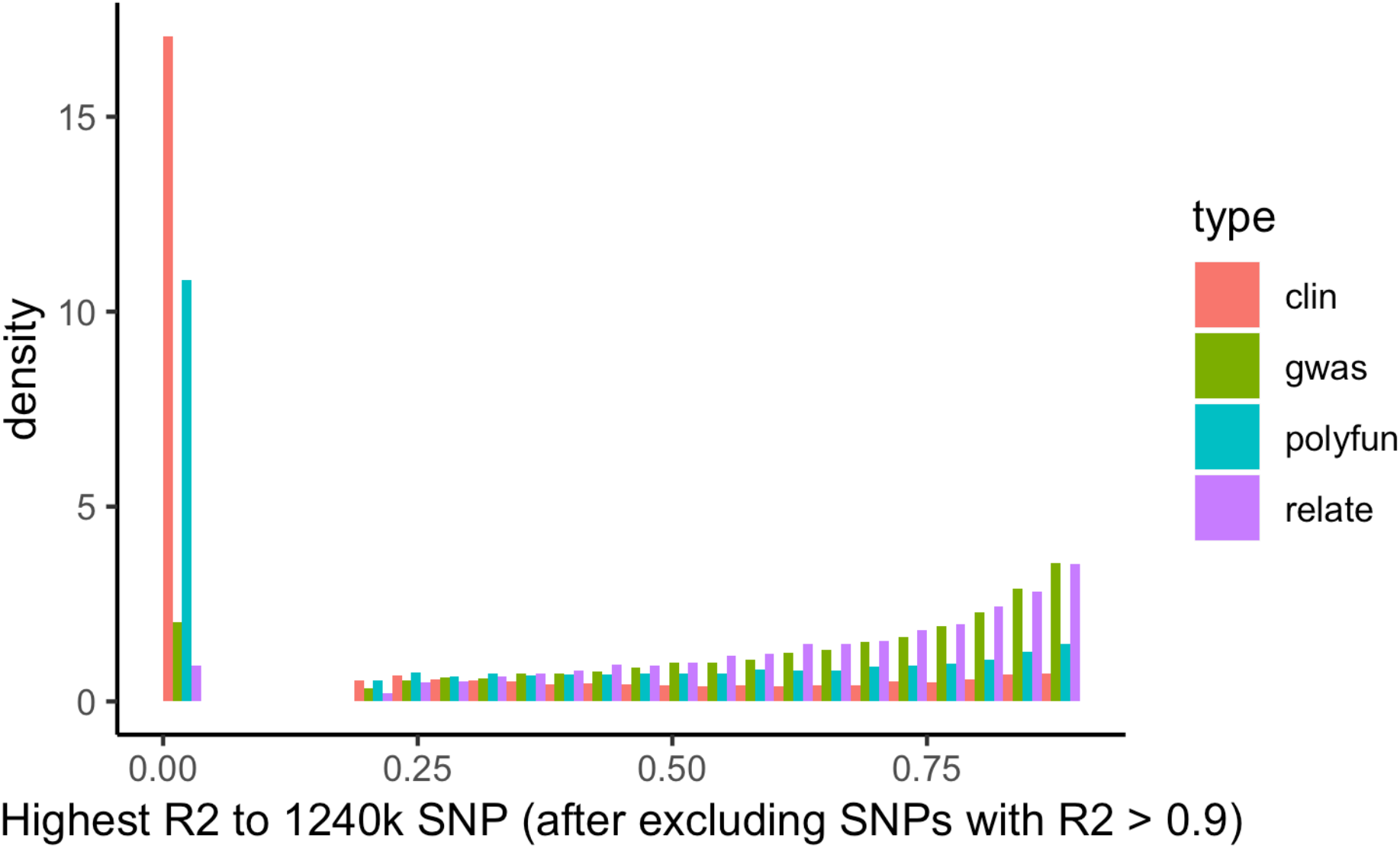
Linkage disequilibrium distribution by source. All SNPs with highest LD<0.2 set to 0.

Finally, we manually added in 15 phenotypically important multi-allelic polymorphisms and 6 insertion/deletion targets where we tiled both alternative alleles (Table S1.3).

**Table S1.3:**
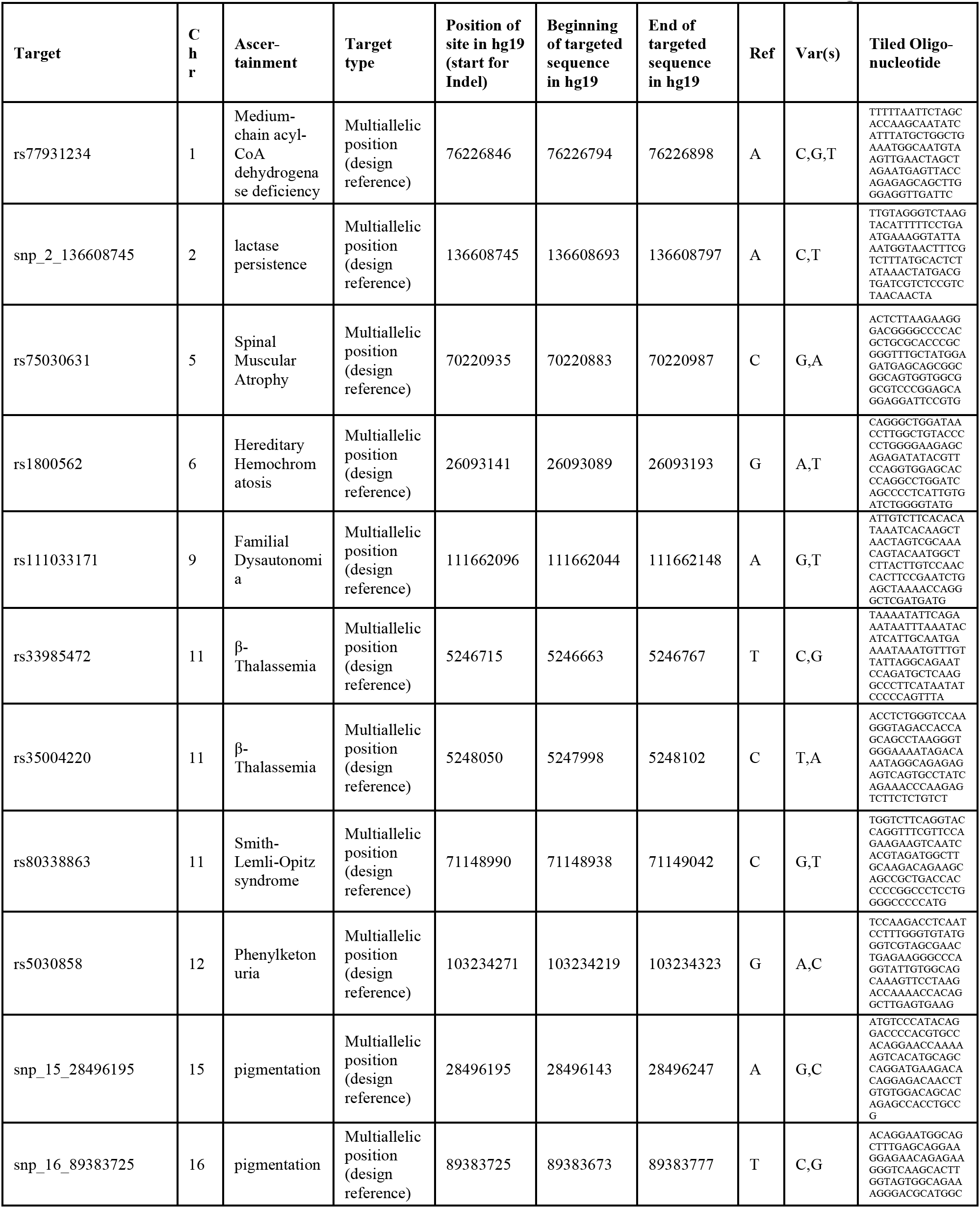
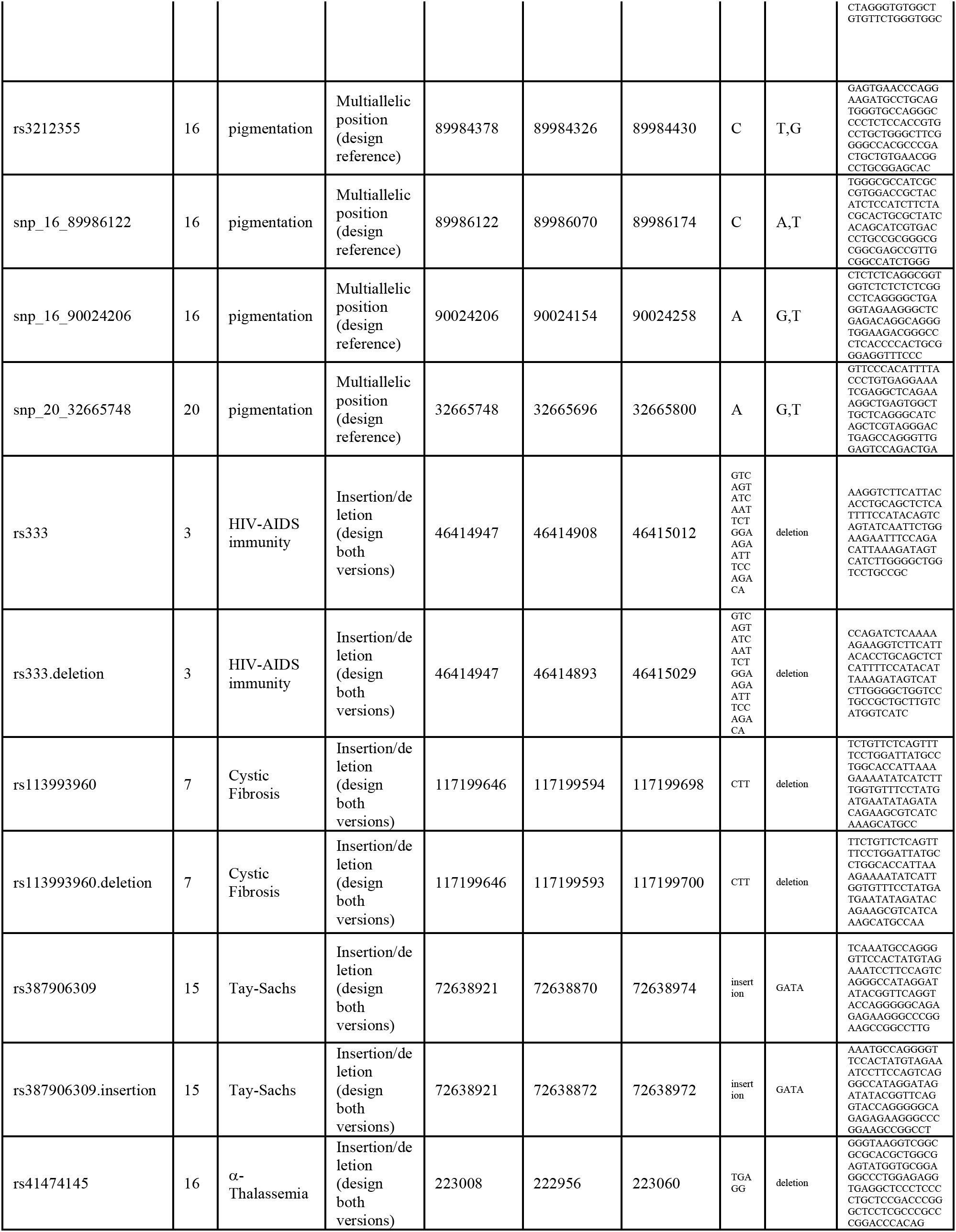
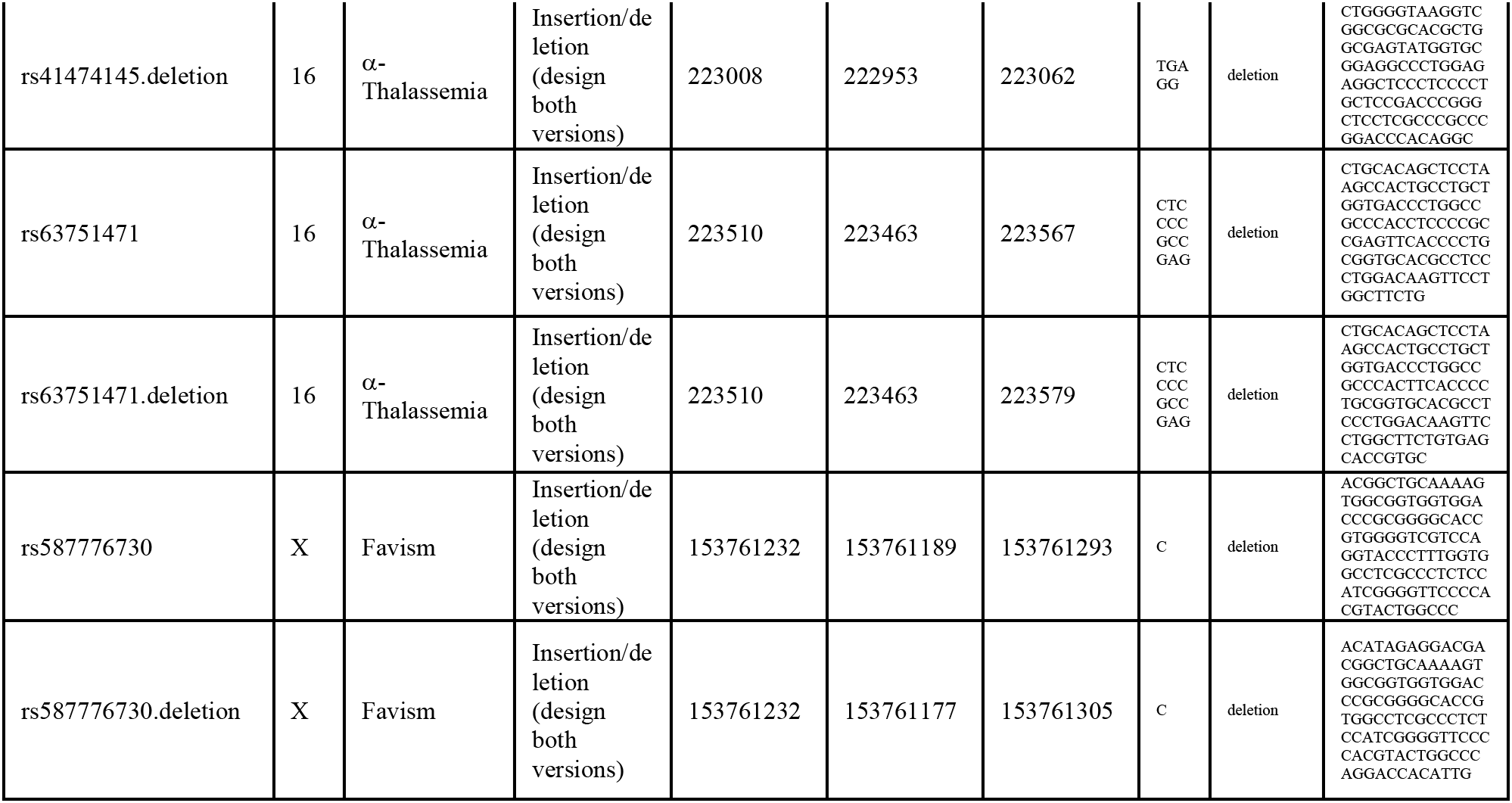
Manual addition of 15 multiallelic SNPs and 6 insertion/deletion targets.

#### (1b) Targeting 81,925 polymorphisms on chromosome Y

To identify Y chromosome targets, we started with 32,670 chromosome Y SNPs from the 1240k reagent. These had been identified by starting with ISOGG 9.77 SNPs (https://isogg.org/), and then merging with SNPs identified as polymorphic in the Simons Genome Diversity Panel (52, 53).

For our redesign, we added in 69,991 Y SNPs from the ISOGG Y SNP index version 14.199 downloaded Nov. 5 (https://isogg.org/). To obtain this list, we started with 88,795 polymorphisms in the download, removed ones with duplicate positions, and restricted to true SNPs that are biallelic for the alleles A/C/G/T.

After merging and removing duplicates, this generated 88,023 SNPs. We reduced this to 81,925 by removing SNPs monomorphic in the existing 1240k enrichment dataset, or that had coverage counts in that dataset of <10%.

In contrast to the 94,586 SNPs identified in Section 1a which represent a supplement to the 1240k content on chromosomes 1-22 and X, for the Y chromosome the 81,925 SNPs we discuss are a replacement of the 1240k content on chromosome Y.

#### (1c) Final count of SNPs

The total number of SNPs targeted for the reagent is:

|  |  |
| --- | --- |
| 1,200,343 | 1240k content on chromosomes 1-22 and Y |
| 94,586 | Newly designed phenotypic discussed in Section 1a |
| 81,925 | Fully redesigned Y chromosome content discussed in Section 1b |
| 1,376,854 | Total |

For each targeted SNP, we randomly selected a third allele to represent each position and flanked it 52bp on either side according to the sequence from the h*g19* reference genome. We then mapped the sequence to hg19. After removing oligonucleotides that mapped unreliably with a score of MAPQ<23, or that mapped to a location that disagreed with the recorded positions, or that was duplicated in its sequence compared to another in the dataset, or that failed other quality controls, our design file targeted 1,352,535 SNPs.

#### (1d) Tiled regions (with either 1x or 2x tiling)

Beyond SNP targeting, we also added in probes to bait additional genomics regions.

- *“Methylation” targets* We are grateful to Steve Horvath and Vagheesh Narasimhan for providing us with the coordinates of 40,000 CpG dinucleotides chosen to be locations where methylation rates are correlated to the skeletally determined ages of ancient individuals. These CpG dinucleotides are also ones where methylation rates have been shown to be well-correlated to the ages of living individuals. Of these targets, we successfully designed single probes for 39,886 (we did not design probes for the others due to repetitive flanking sequence).
- *“Human Accelerated Region (HAR)” targets* We are grateful to Ryan Doan for sharing with us a list of 3,171 Human Accelerated Regions (HARs) spanning 857,339 nucleotides. We tiled each of these regions twice (with 80bp probes overlapping every 40bp).
- *“Gene resequencing” targets* This includes 9 contiguous regions in 3 genes, specified in hg19 coordinates. The segments target SNPs believed to contribute to β-thalessemia (chr. 11: 5247022-5247193 and 5248114-5248429) α-thalessemia (chr. 16: 222873-223052 and 223469-223733), and favism (chr. X: 153220145-153220335, 153760378-153761377, 153761761-153761889, 153763362-153763532, 153764171-153764423, and 153774226-153774316). The SNPs are rs34690599, rs34451549, rs35724775 , rs33915217, rs33971440 , rs33960103 , rs33986703 , rs34716011 , rs63750783 , rs334, rs34598529, rs33944208 , rs111033603, rs281864819 , rs41474145, rs63750404 , rs63751471 , rs33987053 , rs41397847 , rs41464951, rs63751269 , rs137852348, rs137852344, rs72554664, rs72554665, rs72554665, rs137852324, rs137852317, rs137852337, rs2230037, rs137852336, rs137852323, rs137852335, rs137852316, rs137852316, rs137852321, rs137852334, rs137852320, rs137852322, rs2230036, rs387906468, rs137852329, rs137852345, rs137852333, rs137852342, rs5030869, rs587776730, rs76723693, rs137852347, rs137852339, rs137852327, rs74575103, rs137852318, rs137852346, rs137852328, rs137852328, rs137852319, rs137852326, rs137852332, rs137852332, rs137852330, rs5030868, rs267606836, rs5030872, rs5030872, rs137852343, rs137852331, rs137852314, rs2515904, rs137852313, rs137852341, rs1050829, rs137852349, rs1050828, rs137852315, rs76645461, and rs78478128. We tiled segments with 80bp probes staggered every 40bp.

### Supplementary Section 2: EM Algorithm to Correct for Binomial Sampling Variance

The problem we wish to solve is that we have empirical counts of reference and variant alleles for large numbers of known or highly probable heterozygous positions. Here we describe how we deconvolve the noise to learn the underlying distribution of reference bias.

We consider a set of reference and variance counts (typically summing to 100 or more). At SNP *k* we observe *a*_*k*_ reference and *b*_*k*_ variant alleles. We suppose the ‘true’ allele frequency of reference is *z*_*k*_ = *z* which we can think of as the frequency we would observe if the coverage were infinite. We wish to learn the probability distribution of z. We will ignore (in this note) the case that the observed counts are not polymorphic, so we assume *a*_*k*_, *b*_*k*_ ≥1.

Let us model *z*_*k*_ as lying on a mesh; for instance, *z*_*k*_ = *i*/100 for some i = 1…99. We propose to estimate *p*_*i*_ = (*z*_*k*_ = *i*/100). Write *α*_*i*_ = i/100; *β*_*i*_ = (100-i)/100. We see that the log likelihood of our observation for SNP k is:

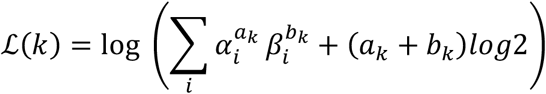

The last term is not essential, but good technique is to score against some random model; here that a_k_ is from tossing a fair coin toss (50% probability heads). The overall log likelihood is:

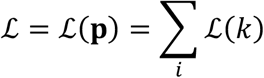

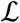 is easily maximized by an EM algorithm. Write:

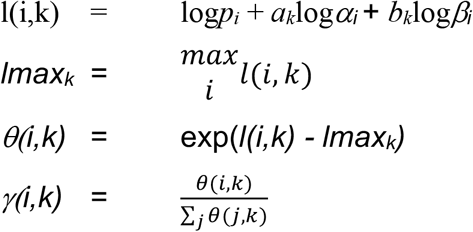

Thus, *γ(i,k)* is the posterior probability that *z*_*k*_ = *α*_*i*_. Reestimates are now simply:

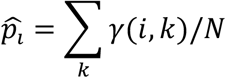

where *N* is the number of SNPs. Standard EM shows that:

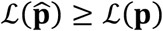

We iterate until convergence. We implemented this in C to produce the inferences in Figure 5.

## Notes

https://www.dropbox.com/sh/h024odwt5w1yc37/AAC9jCMhhOncXQRBaWMWOzPla?dl=0

### Competing Interest Statement

The authors have declared no competing interest.

